# A global pangenome for the wheat fungal pathogen *Pyrenophora tritici-repentis* and prediction of effector protein structural homology

**DOI:** 10.1101/2022.03.07.482416

**Authors:** Paula Moolhuijzen, Pao Theen See, Gongjun Shi, Harold R. Powell, James Cockram, Lise N. Jørgensen, Hamida Benslimane, Stephen E. Strelkov, Judith Turner, Zhaohui Liu, Caroline S. Moffat

**Affiliations:** Centre for Crop Disease and Management, School of Molecular and Life Sciences, Curtin University, Bentley, Western Australia, Australia; Department of Plant Pathology, North Dakota State University, Fargo, North Dakota, USA; Department of Life Sciences, Centre for Integrative Systems Biology and Bioinformatics, Imperial College London, London, England, United Kingdom; NIAB, 93 Lawrence Weaver Road, Cambridge, CB3 0LE, United Kingdom; Department of Agroecology, Aarhus University, Slagelse, Denmark; Département de Botanique, Ecole Nationale Supérieure Agronomique (ENSA), Hassan Badi, El-Harrach, Algiers, Algeria; Department of Agricultural, Food and Nutritional Science, University of Alberta, Edmonton, AB, Canada; Fera Science Ltd., York, YO41 1LZ, United Kingdom

**Keywords:** Necrotrophic fungi, plant pathogen, toxin

## Abstract

The adaptive potential of plant fungal pathogens is largely governed by the gene content of a species, comprised of core and ancillary genes across the pathogen isolate repertoire. To approximate the complete gene repertoire of a globally significant crop fungal pathogen, a pan genomic analysis was undertaken for *Pyrenophora tritici-repentis* (Ptr), the causal agent of tan (or yellow) spot disease in wheat.

In this study, fifteen new Ptr genomes were sequenced, assembled and annotated, including isolates from three races not previously sequenced. Together with eleven previously published Ptr genomes, a pangenome for twenty-six Ptr isolates from Australia, Europe, North Africa and America, representing nearly all known races, revealed a conserved core-gene content of 57% and presents a new Ptr resource for searching natural homologues using remote protein structural homology. Here, we identify for the first time a nonsynonymous mutation in the Ptr effector gene *ToxB*, multiple copies of *toxb*, a distant natural *Pyrenophora* homologue of a known *Parastagonopora nodorum* effector, and clear genomic break points for the *ToxA* effector horizontal transfer region.

This comprehensive genomic analysis of Ptr races includes nine isolates sequenced via long read technologies. Accordingly, these resources provide a more complete representation of the species, and serve as a resource to monitor variations potentially involved in pathogenicity.

**Author Notes:** All supporting data, code and protocols have been provided within the article or through supplementary data files. Five supplementary data files and fifteen supplementary figures are available with the online version of this article.

**Impact Statement:** Our *Pyrenophora tritici-repentis* (Ptr) pangenome study provides resources and analyses for the identification of pathogen virulence factors, of high importance to microbial research. Key findings include: 1) Analysis of eleven new sequenced (with three new races not previously available) and previously published isolates, 26 genomes in total, representing the near complete Ptr race set for known effector production collected from Australia, Europe, North Africa and the Americas. 2) We show that although Ptr has low core gene conservation, the whole genome divergence of other wheat pathogens was greater. 3) The new PacBio sequenced genomes provide unambiguous genomic break points for the large *ToxA* effector horizontal transfer region, which is only present in ToxA producing races. 4) A new web-based Ptr resource for searching *in silico* remote protein structural homology is presented, and a distant natural *Pyrenophora* protein homologue of a known effector from another wheat pathogen is identified for the first time.

**Data Summary:** The sources and genomic sequences used throughout this study have been deposited in the National Centre for Biotechnology Information (NCBI), under the assembly accession numbers provided in Tables 1 and 2 (available in the online version of this article). The new M4 resource for protein structural homology is freely available through the BackPhyre web-portal URL, http://www.sbg.bio.ic.ac.uk/phyre2/.

## Introduction

Tan (or yellow) spot, caused by the necrotrophic fungal pathogen *Pyrenophora tritici-repentis* [(Died.) Drechs.] (abbreviated to Ptr), can occur on both bread wheat (*Triticum aestivum* L.) and durum wheat (*T. turgidum* subsp. *durum* L.). A globally significant disease of economic importance (Murray GM and Brennan JP 2009, Benslimane, Lamari et al. 2011), tan spot can reduce crop production with up to 31% yield losses reported (Bhathal, Loughman et al. 2003).

During infection, necrotrophic fungi secrete necrotrophic effectors (NEs) that interact with the corresponding sensitivity genes in the host wheat lines (Friesen, Zhang et al. 2008, Ciuffetti, Manning et al. 2010, Faris, Liu et al. 2013, Downie, Lin et al. 2021). To date, Ptr has three known NEs, Ptr ToxA, Ptr ToxB and Ptr ToxC that produce either necrosis or chlorosis symptoms on their sensitive wheat genotypes (Ciuffetti, Tuori et al. 1997, Strelkov, Lamari et al. 1999). It is the different combinations (and absence) of these NEs that have been used to define different Ptr races (Lamari and Strelkov 2010, Faris, Liu et al. 2013), including race 1 (Ptr ToxA and Ptr ToxC), race 2 (Ptr ToxA), race 3 (Ptr ToxC), race 4 (no Ptr ToxA, Ptr ToxB or Ptr ToxC), race 5 (Ptr ToxB), race 6 (Ptr ToxB and Ptr ToxC), race 7 (Ptr ToxA and Ptr ToxB) and race 8 (Ptr ToxA, Ptr ToxB and Ptr ToxC). However, there are reports of isolates beyond the current classification (Ali S, Gurung S et al. 2010, Benslimane, Lamari et al. 2011, Kamel, Cherif et al. 2019). AR CrossB10 from North Dakota, USA, was such an isolate that produces both Ptr ToxC with an unknown effector, which has been recently sequenced (Kariyawasam, Wyatt et al. 2021) and will subsequently be referred to here as “race unknown” (Moolhuijzen, See et al. 2018). In addition to the three known effectors, the presence of novel NEs has been suggested in several studies (Tuori, Wolpert et al. 1995, Andrie, Pandelova et al. 2007, Ali S, Gurung S et al. 2010, Rybak, See et al. 2017, See, Marathamuthu et al. 2018). These findings make the sequencing of new isolates a priority to capture and understand the complete gene repertoire for Ptr.

Genome sequencing projects for fungal pathogens using single molecule long reads, such as PacBio and Oxford Nanopore technologies, have significantly improved our understanding of pathogen genomes, as they allow near complete genome assembly. In particular, the wheat fungal pathogens *Fusarium graminearum* (cause of fusarium head blight), Ptr, *Parastagonospora nodrum* (Sn, cause of Septoria nodorum blotch) and *Zymoseptoria tritici* (Zt, cause of Septoria tritici blotch) are known to have highly variable genomes characterised by gene loss and duplication events as well as large-scale genome rearrangements (Manning, Pandelova et al. 2013, Richards, Wyatt et al. 2017, Moolhuijzen, See et al. 2018, Badet, Oggenfuss et al. 2020, Alouane, Rimbert et al. 2021, Bertazzoni, Jones et al. 2021). To understand the genome composition of a species, the protein-coding genes from all available isolates are clustered based on the sequence identity of conserved protein domains into core (genes shared by all isolates) and ancillary/accessory (genes absent in one or more isolates) groups. The union of the core and accessory groups for the collection of isolates is then referred to as the pangenome, which is larger than the genome of any one individual (Vernikos, Medini et al. 2015). Depending on the number of sequenced isolates, associations to distinct habitats and phenotypes may then be detected within a pathogen species (Vernikos, Medini et al. 2015).

In this study, fifteen new Ptr isolates collected from Europe (Denmark, Germany and the United Kingdom), North Africa (Algeria and Tunisia) and the Americas (Brazil, Canada and the USA) were **s**equenced, assembled and annotated, for comparative analysis with eleven previously published Australian and North American Ptr isolates (Manning, Pandelova et al. 2013, Moolhuijzen, See et al. 2018, Moolhuijzen, See et al. 2019). A total of 26 annotated Ptr genomes, which represent nearly all known Ptr races, are presented here for a pangenome analysis to determine whole genome phylogeny and sequence variations in relation to core and ancillary genes. Ptr proteins are then further explored *in silico* to identify remote natural structural homology between different necrotrophic fungal species (not acquired by a horizontal gene transfer).

## Materials and methods

### Isolate collection and DNA extraction

Ptr isolates were collected from Algeria (Alg130 and Alg215), Brazil (Biotrigo9-1), Canada (90-2), Denmark (EW306-2-1, EW4-4, and EW7m1), Germany (SN001A, SN001C and SN002B), USA (86-124 and Ls13-192), United Kingdom (CC142) and Tunisia (T199 and T205). All isolates were collected from bread wheat (*T. aestivum* L.), except Alg215 which was collected from durum wheat (*T. turgidum* subsp. *durum* L.). Fungi were grown on V8-PDA agar according to (Moffat, See et al. 2014). Genomic DNA was extracted using a BioSprint 15 DNA Plant Kit (Qiagen, Hilden, Germany) with some modifications. Briefly, DNA was extracted using the BioSprint 15 automated workstation, according to the manufacturer’s instructions, from 3-day old mycelia grown in Fries 3 medium (Moffat, See et al. 2014). DNA was further treated with 50 µg/ml of RNase enzyme (Qiagen, Hilden, Germany) for 1 h followed by phenol/chloroform extraction, precipitation with sodium acetate and ethanol, and finally resuspension in Tris-EDTA buffer.

### Isolate pathotyping

Ptr isolates were pathotyped for race classification through infection assays of differential wheat genotypes differing in their specific effector sensitivities. The wheat genotypes used were Glenlea (Ptr ToxA-sensitive), 6B662 (Ptr ToxB-sensitive), 6B365 (Ptr ToxC-sensitive) and Auburn or Salamouni (insensitive to all three effectors).

Two-week-old wheat (*T. aestivum* L.) seedlings were inoculated by spraying conidia onto the whole plants evenly at a rate of 3,000 conidia/ml and grown at 20 °C under a 12-h day/night cycle in a controlled growth chamber (Moffat, See et al. 2014). The second leaves were harvested 7-days post-inoculation, visually inspected for symptoms (Lamari, Sayoud et al. 1995) and photographed. The inoculation experiments were repeated twice with three replicate plants per wheat genotype.

### Ptr isolate sequencing and genome assembly

Genomic DNA from four Ptr isolates was sequenced using the PacBio Sequel system, 90-2 (Novogene, China), Biotrigo9-1 (Novogene, USA), Ls13-192 and 86-124 (Mayo Clinic, Minnesota, USA). Error correction and *de novo* genome assembly of PacBio reads was completed with Canu version v2.1.1 (Koren, Walenz et al. 2017) with the following options (genomeSize=43, useGrid=TRUE, maxThreads=28, merylThreads=28, ovlThreads=28 ovlMerThreshold=500 and gridOptionsOBTOVL=“--cpus-per-task=28) on computer resources (Broadwell Intel Xeon cores, 100 Gb/s Omni-Path interconnect and 128GB of memory per compute node) at Pawsey Supercomputing Centre, Perth, Western Australia. Previously generated Illumina 150 bp paired end DNA sequence reads of 86-124 genomic DNA (Moolhuijzen, See et al. 2018) and Biotrigo9-1 Illumina sequence (this study) were aligned to the contigs using BWA V0.7.17-r1188 (Li H and Durbin R 2009), and the sorted alignment bam files then used for further base error corrections using Pilon v1.24 (Walker, Abeel et al. 2014).

The genomic DNA for an additional 11 Ptr isolates (EW306-2-1, EW4-4, and EW7m1, SN001A, SN001C, SN002B, CC142, Alg130, Alg215, T199 and T205) was sequenced using Illumina Hi-Seq 150 bp pair-end reads by the Australian Genome Research Facility (AGRF). Isolate sequence data was quality checked with FASTQC (Andrews 2011), trimmed for poor quality, ambiguous bases and adapters using Skewer (Jiang, Lei et al. 2014) and Trimmomatic v0.22 (Bolger, Lohse et al. 2014) with a read head crop of 6 bp and minimum length of 100 bp. *De novo* genome assembly was undertaken using SPAdes version v3.10.0 (Bankevich, Nurk et al. 2012).

### Gene prediction and functional annotation

Ptr sequenced genomes were soft masked for low complexity, as well as known transposable elements, using RepeatMasker (RM) (Chen 2004) v. open-4.0.6 with rmblastn version 2.2.27+ on RepBase (Kohany O, Gentles AJ et al. 2006) RM database version 20150807 (taxon=fungi). *Ab-initio* gene predictions were made with GeneMark-ES v4.33 (--ES --fungus --cores 16) (Borodovsky and Lomsadze 2011) and CodingQuarry v1.2 Pathogen Mode (PM) (Testa, Hane et al. 2015), assisted by RNA-Seq (Moolhuijzen, See et al. 2018) genome alignments using TopHat2 (Kim and Salzberg 2011) for a minimum intron size of 10 bp. The Ptr M4 and Pt-1C-BFP reference proteins (Manning, Pandelova et al. 2013, Moolhuijzen, See et al. 2018) were aligned using Exonerate v2.2.0 (--minintron 10 --maxintron 3000) protein2genome mode (Slater and Birney 2005). Gene annotations were assigned from BLASTX (v2.2.26) (Shiryev, Papadopoulos et al. 2007) searches against Uniref90 (Oct 13, 2020), NCBI Refseq (taxon=Pezizomycotina) (Oct 13, 2020) and InterProScan v5.17-56 (Quevillon, Silventoinen et al. 2005) protein databases. Sequence domains were assigned by RPS-BLAST (v2.2.26) against Pfam v33.1, Smart v6.0 and CDD v3.19 databases. The blast protein and domain searches were then summarised using AutoFACT v3.4 (Koski LB, Gray MW et al. 2005).

Proteins were screened for a signal peptide using SignalP v5.0b (Petersen TN, Brunak S et al. 2011). Effector predictions were conducted on proteins with signal peptides using EffectorP v3.0 (Sperschneider, Dodds et al. 2018, Sperschneider and Dodds 2021). To ensure the same prediction methods were used for comparative analyses, SignalP V5.0b and EffectorP v3.0 (Sperschneider, Dodds et al. 2018, Sperschneider and Dodds 2021) were used to update the effector gene predictions on all the publicly available isolate genomes (Supplementary data 1). All predicted proteins were also ranked using Predector v1.1 (Jones, Rozano et al. 2021) (Supplementary data 1). Gene completeness was accessed using BUSCO v3, lineage fungi (Seppey, Manni et al. 2019).

### Comparative genomics

To conduct comparative analyses across the Class Ascomycota, publicly available isolate genomes were downloaded from the National Centre for Biotechnology Information (NCBI) GenBank. These included *Bipolaris* (*B. cookei*, *B. maydis*, *B. sorokiniana*, *B. zeicola*), *Leptosphaeria* (*L*. *maculans*)*, Parastagonospora* (*P. nodorum*), *Pyrenophora* (*Pyrenophora teres* f*. teres, Pyrenophora teres* f*. maculata*, *Pyrenophora serminiperda*) and *Zymoseptoria* (Z. *tritici*) (Syme, Tan et al. 2016, McDonald, Ahren et al. 2017, Richards, Wyatt et al. 2017, Syme, Martin et al. 2018, Badet, Oggenfuss et al. 2020) (Supplementary data 2). The published genomes of *P. tritici-repentis* isolates Pt-1C-BFP, DW5, DW7, SD20 (Manning, Pandelova et al. 2013), Ptr134, Ptr239, Ptr11137, Ptr5213, M4, 86-124 (Moolhuijzen, See et al. 2018), AR CrossB10 (Kariyawasam, Wyatt et al. 2021) and V1 (Moolhuijzen, See et al. 2019) were also included for analysis.

Genome nucleotide pairwise distance was calculated with Phylonium v1.5 (Klotzl and Haubold 2020) with two-pass enabled and 100 bootstrap matrices. Whole genome phylogenetic trees were constructed using Phylip 1:3.695-1 (Retief 2000), consensus program v3.695 on 100 Kitsch and neighbour joining v3.695 trees. The tree was then visualised using FigTree v1.4.4. Genomic nucleotide regions were compared between isolates using NUCmer v3.1 (Delcher, Salzberg et al. 2003) and Easyfig v 2.2.3 (Sullivan, Petty et al. 2011).

To determine the presence, copy number and percent identity of all genes in Ptr, the gene nucleotide sequences from all 26 isolates were aligned to all 26 genomes using GMAP version 2021-05-27 with options “-f 2 -t 48 -n 300 --max-intronlength-middle=1000 --max-intronlength-ends=1000 --fulllength --trim-end-exons=0 --alt-start-codons --canonical-mode=1 -- --max-deletionlength=20”. Isolate mRNA Pearson correlations and predicted effector protein lengths and scores were analysed using R v4.0.3 (Team” 2021) using the R packages corrplot v0.84, ggplot2 v3.3.3, ggridges v0.5.3 and pheatmap v1.0.12. The analysis and data are available in v1.3.1093 RStudio (RStudio-Team 2020) markdown notebook https://github.com/ccdmb/PTR-60.

Isolate reads were aligned to the isolate M4 reference genome using BWA 0.7.14-r1138 and coverage (10 kb windows) was calculated using BedTools (genomecov) v2.17.0 on SamTools v 0.1.19-96b5f2294a sorted bam files. Regions of absence were then plotted using Circos v0.69-3 and R v3.5.1, bioconductor package chromPlot v1.10.0.

### Protein orthologous clustering and effector analysis

Predicted protein data for all available Ptr isolates were clustered using OrthoFinder v2.5.2 (Emms and Kelly 2015). The predicted effector groups (with signal peptides) were then screened for three-dimensional (3-D) protein model predictions using the Protein Homology/analogY Recognition Engine V 2.0 Phyre2 (Kelley, Mezulis et al. 2015) batch processing mode. The predicted models were superimposed on the best ranked template to find the largest subset of atoms within an approximate threshold of 3.5 Å, which was adjusted based on the size of the aligned proteins using iMol (Rotkiewicz 2007). Protein sequences with high confidence (Phyre^2^ ≥ 90%) predicted 3-D protein models were also searched against the Plant Host Interactions database (PHI-base) of known pathogenic phenotypes (Urban, Cuzick et al. 2017), at an expected value threshold of ≤ 1e-10 for significant alignments. Hidden Markov Model (HMM) libraries were created for the whole genome of Ptr isolate M4, which has been made publicly available through the online resource BackPhyre, Imperial College, London (Kelley, Mezulis et al. 2015).

## Results

### PacBio genome sequencing, assembly and annotation of four *P. tritici repentis* isolates

A total of four Ptr genomes comprising two race 4 (lack all three known Ptr effectors) isolates (North Dakota (USA) isolate Ls13-192 (Guo, Shi et al. 2020) and Canadian isolate 90-2 (Lamari, Gilbert et al. 1998)), and two race 2 (producing Ptr ToxA only) isolates (Brazilian isolate Biotrigo9-1 (Bertagnolli, Ferreira et al. 2019) and Canadian isolate 86-124 (Lamari and Bernier 1989)) were sequenced using PacBio technology, assembled and protein-coding genes were predicted for comparative analysis.

The assembled Ptr genomes ranged in size from 37.56 Mb to 42.19 Mb (Table 1) and of these, the known effector producing isolates (86-124 and Biotrigo9-1) had a size comparable to previously PacBio sequenced genomes (M4 and DW5) (Moolhuijzen, See et al. 2018, Moolhuijzen, See et al. 2020). The race 4 isolate not producing known effectors, Ls13-192, had the smallest genome size at 37.56 Mb, at least 2 Mb smaller than all the known effector-producing isolate genomes, but similar to Pt-1C-BFP which was sequenced prior to the availability of third generation long read technologies and which lacks some representation of repeat/complex genomic regions (Manning, Pandelova et al. 2013). Our four new assemblies were more fragmented than the previously assembled genomes M4 and DW5 (Moolhuijzen, See et al. 2018, Moolhuijzen, See et al. 2020). In particular race 4 isolate 90-2 was fragmented into 162 contigs, over twice as many contigs as compared to race 2 isolate Biotrigo9-1 and race 4 isolate Ls13-192. The four genome assemblies had a BUSCO quantitative assessment greater than 98.9% for completeness with respect to gene content (Supplementary Fig. S1).

**Table 1.**
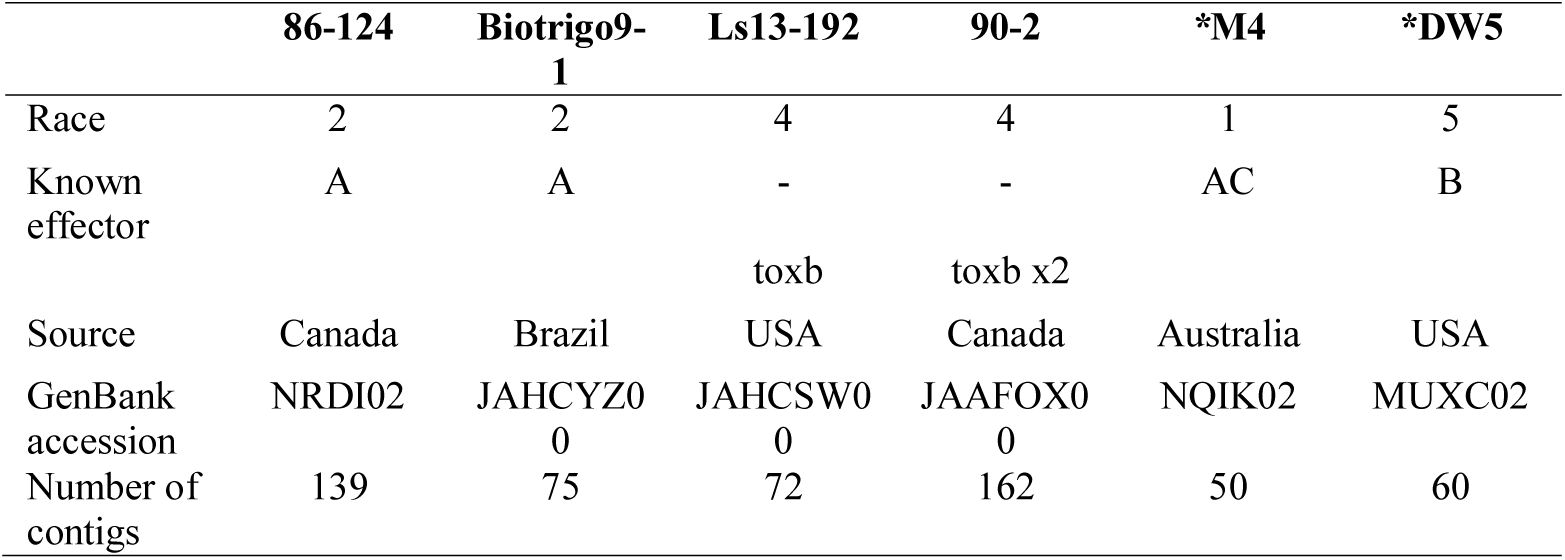
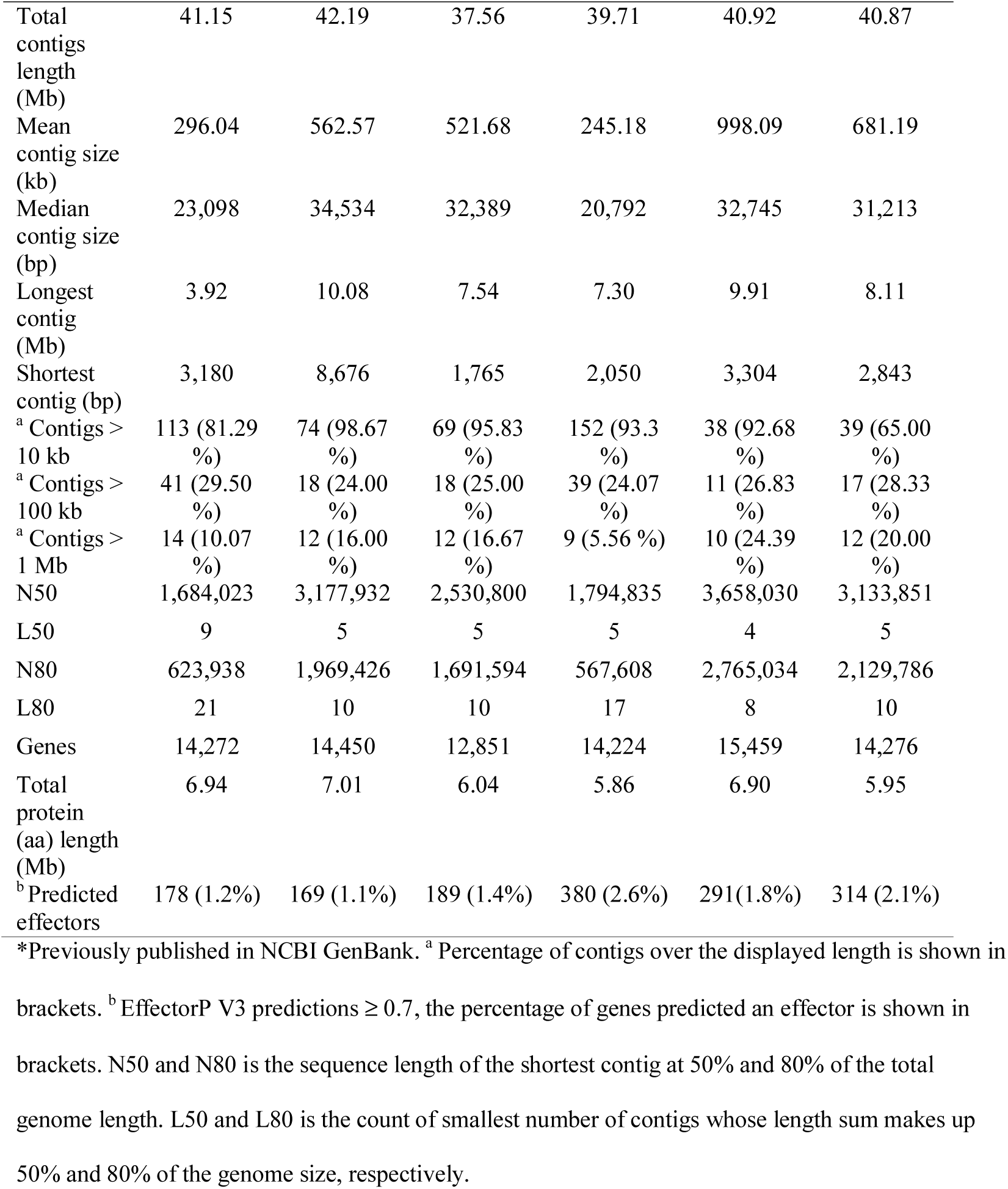
Summary statistics for our four PacBio sequenced Ptr genome assemblies, compared with those of two previously published Ptr assemblies.

The number of predicted protein-encoding genes for our new PacBio sequenced genome assemblies ranged from 12,851(Ls13-192) to 14,450 (Biotrigo9-1) (Table 1). The number of predicted protein effectors for race 2 isolates 86-124 and Biotrigo9-1 and race 4 isolate Ls13-192 was lower than the numbers predicted for race 1 isolate M4 and race 5 isolate DW5. However, race 4 isolate 90-2 had the highest number of proteins predicted as effectors, due to a higher gene copy number identified later in the protein clustering analysis. Furthermore, in the race 4 isolates a single *toxb* (found in non-pathogenic Ptr isolates and having no toxic activity (Kim and Strelkov 2007, Figueroa Betts, Manning et al. 2011) was detected in Ls13-192 on contig 4 (113,627-113,893 bp) and an exact *toxb* duplication event was detected in 90-2 on contig 37 (termed here *toxb1*, 15,199-15,465 bp) and on contig 42 (termed *toxb2*, 15,135-15,401 bp). The *toxb* genes appeared close to a contig end. Ls13-192 contig 4 and 90-2 contigs 37 and 42 have contig assembly sizes 3,110,128 Mb, 116 kb and 87 kb, respectively. No *toxb* gene coding region, protein or nucleotide sequence variations were identified (Supplementary Fig. S2 and S3). *ToxA* was identified in race 2 isolates 86-124 (contig 17, 764,135-764,722 bp) and Biotrigo9-1 (contig 7, 1,370,173-1,370,760 bp), no gene coding region, nucleotide or protein sequence variations were found. The Ptr-specific hairpin element (PtrHp1) *ToxA* 3’UTR insertion previously identified (Moolhuijzen, See et al. 2018) was not detected in the *ToxA* 3’UTR region of these genomes.

The four new assembled and annotated genomes Ls13-192, 86-124, 90-2 and Biotrigo9-1 have been deposited in NCBI GenBank and can be found under accession numbers JAHCSW000000000, NRDI02000000, JAAFOX000000000 and JAHCYZ000000000, respectively.

### Illumina genome sequencing, assembly and annotation of eleven *P. tritici repentis* isolates

Whole genome Illumina sequencing and assembly was then undertaken for eleven new Ptr genomes comprised of isolates from Denmark (EW306-2-1, EW4-4 and EW7m1), Germany (SN001A, SN001C and SN002B), United Kingdom (CC142), Algeria (Alg130 and Alg215) and Tunisia (T199 and T205). The assembled Ptr genomes ranged in size from 34.15 Mb to 35.18 Mb (Table 2), comparable to previous Illumina Ptr isolate assembly sizes (Moolhuijzen, See et al. 2018).

**Table 2.**
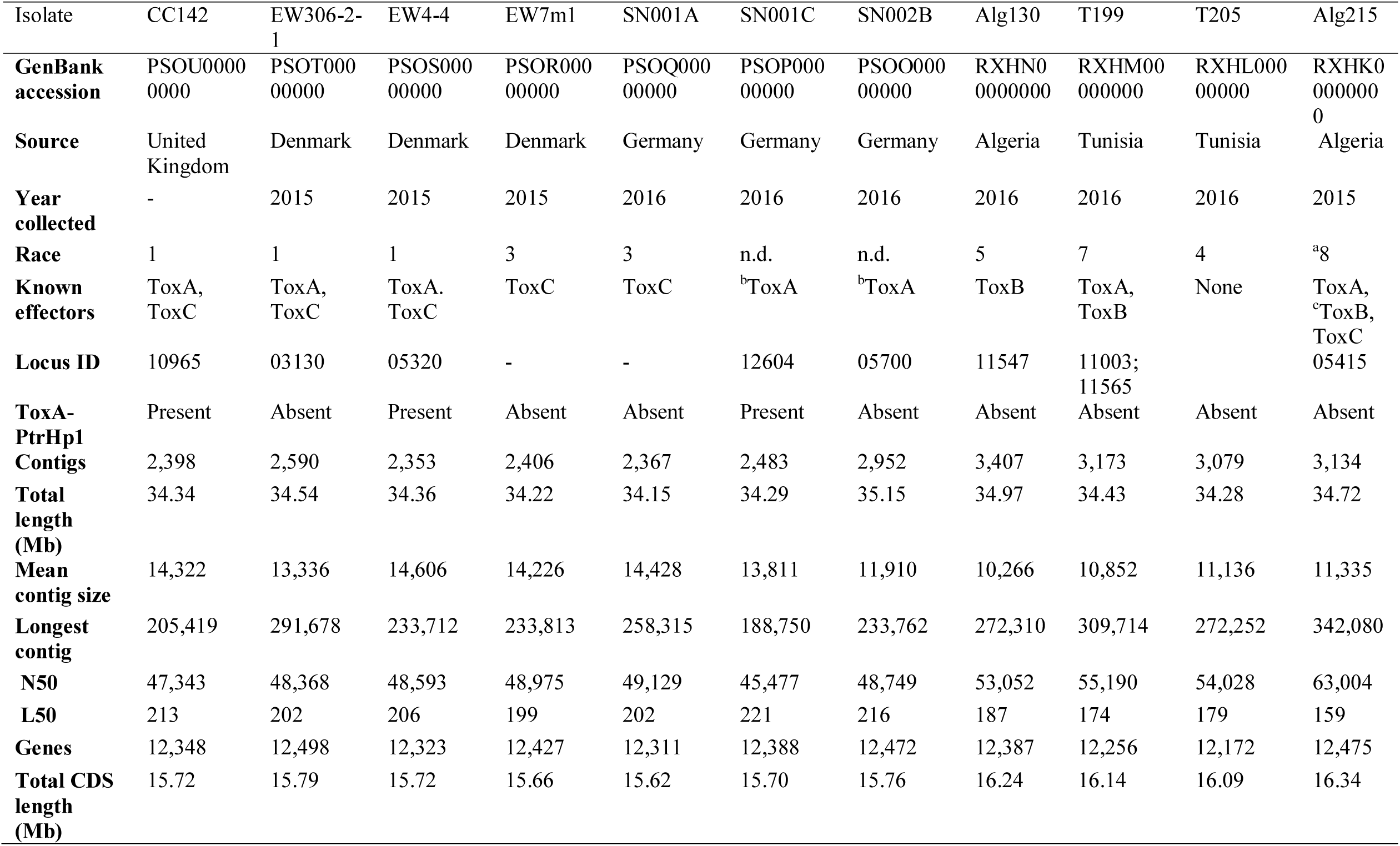
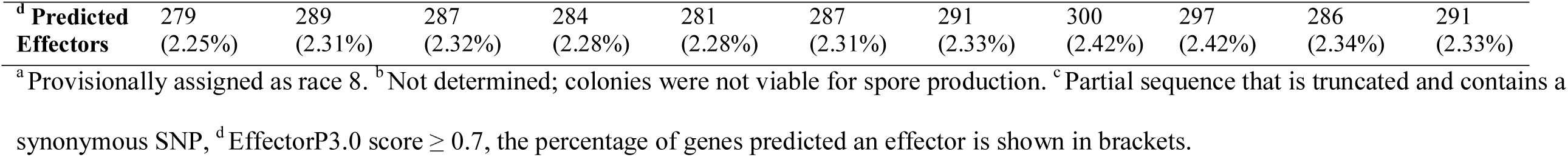
Illumina sequenced genome assemblies of 11 new Ptr isolates. Table shows isolate source, race and *de novo* assembly statistics.

The number of predicted protein-encoding genes ranged between 12,172 to 2,498 for the assembled genomes. Of these, 279 to 300 effectors were predicted with a probability score ≥ 0.7. *ToxA* was identified in isolates T199, Alg215, CC142, EW3061-2-1, EW4-4, SN001C and SN001B, and *ToxB* was identified in the Alg130 genome. *ToxA* and *ToxB* were both detected in T199 and Alg215 genomes, however, the Alg215 *ToxB* sequence was partial (due to a sequence inversion in the 5’ end of the gene), truncated by 33 amino acid residues in the protein N-terminus which includes the encoded signal peptide (amino acid positions 1 to 22) (Supplementary Fig. S4). Furthermore, a single nonsynonymous substitution (I > R, residue position 17) was detected. Neither *ToxA* or *ToxB* were detected in isolates EW7m1, SN001A and T205.

The Ptr-specific hairpin element (PtrHp1) *ToxA* 3’ UTR insertion previously identified in isolates EW306-2-1 and EW4-4 (Moolhuijzen, See et al. 2018) was also detected in *ToxA* 3’ UTR for our United Kingdom isolate CC142, but not in the remaining North African *ToxA* isolates T199 and Alg215 (Table 2).

The plant infection assays on the wheat differential lines confirmed CC142, EW306-2-1 and EW4-4 as race 1 isolates (producing ToxA and ToxC), EW7m1 and SN001A as race 3 isolates (producing ToxC), Alg130 as a race 5 isolate (producing ToxB), T199 as a race 7 isolate (producing ToxA and ToxB) and T205 as a race 4 isolate (no ToxA, ToxB or ToxC production) (Supplementary Fig. S5 and S6). Due to the truncated *ToxB* gene in isolate Alg215 and a weaker chlorosis phenotype on the ToxB wheat differential lines, Alg215 has been provisionally classified as a race 8 isolate (producing ToxA, ToxB and ToxC) (Table 2). The SN001C and SN002B isolates could not be tested for race classification because the colonies sporulated poorly; nonetheless, *ToxA* was present and *ToxB* was absent in the genome sequence for both isolates. As ToxC production in SN001C and SN002B remains unknown, they could be race 1 or 2.

All the assembled and annotated genomes have been deposited in NCBI GenBank and can be found under accession numbers PSOO00000000-PSOU00000000 and RXHK00000000-RXHN00000000.

### Whole genome comparative analyses

Whole genome phylogenetic analysis of the 26 Ptr isolates, sourced from the major wheat growing regions in the Americas, Australia, Europe and North Africa (Fig. 1A), showed distinct clades for European and North African geographic locations (Fig. 1B). Surprisingly, isolate Alg215 from North Africa did not cluster with the remaining North African isolates. On genome alignment to the reference genome of isolate M4, a large 1 Mb distal region on M4 contig 1 and many smaller regions were absent in Alg130, T199 and T205 but present in Alg215 (Supplementary Fig. S7). Furthermore, branches for race 4 (that do not produce ToxA, ToxB or ToxC) isolates (SD20, 90-2 and Ls13-192) had the greatest phylogenetic distances from the known effector producing isolate groups, while race 4 T205 and SD20 (both Illumina sequence) did not cluster. In particular, isolates SD20 (USA) and 90-2 (Canada) were more distant than the isolate Ls13-192 (USA).

**Fig. 1.**
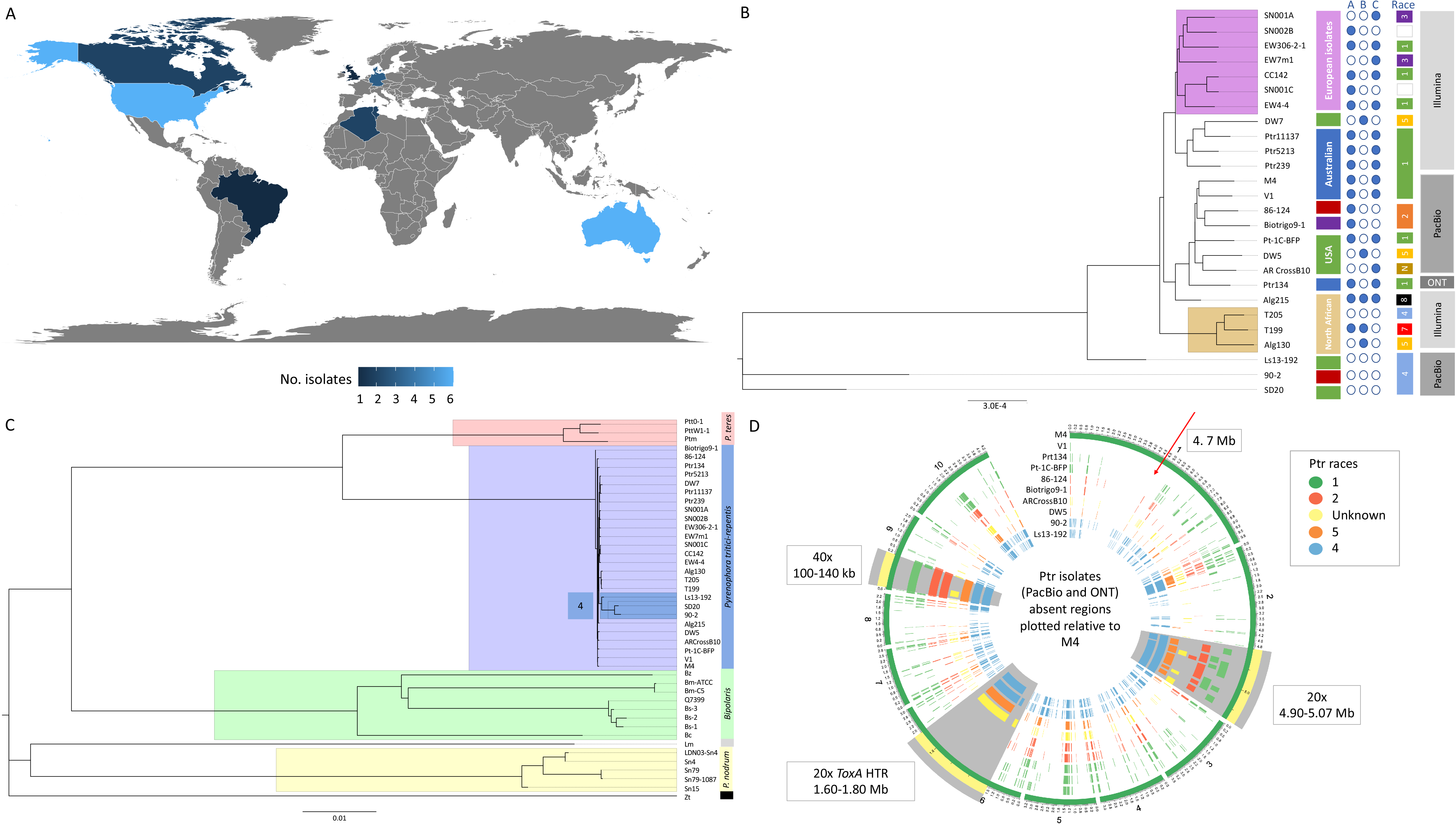
Whole genome analysis of Ptr isolates. A) Geographic source and number of Ptr isolates currently available and analysed. Legend shows the number of isolates. B) Whole genome phylogenetic tree of Ptr isolates from Illumina sequencing (Alg130, T199, T205, Alg215, CC142, EW306-2-1, EW4-4, EW7m1, DW7, Pt-1C-BFP, Ptr239, Ptr11137, Ptr5213, SD20, SN001A, SN001C, SN002B), PacBio Technologies (86-124, 90-2, AR CrossB10, Biotrigo9-1, DW5, Ls13-192, M4 and V1) and Oxford Nanopore Technologies (Ptr134). The unrooted neighbour joining phylogenetic tree displays clades for the European (violet) and North Africa (tan) isolates. Geographic source of the other isolates are Australian (blue), USA (green), Canada (red) and Brazil (purple). The race 4 isolates (Ls13-192, 90-2 and SD20) have the greatest distance from the clade of known effector producing isolates. C) Unrooted Neighbour joining phylogenetic tree for Ptr (purple clade), *Pyrenophora teres* (*P. teres* f. *maculata* (Ptm) and *P. teres* f. *teres* (Ptt)) (orange clade), *Bipolaris* (*B. sorokiniana* (Bs1-3 and Q7399), *B. maydis* (Bm-ATCC and Bm-C5) and *Bipolaris zeicola* (Bz) (green clade), *Parastagonospora nodorum* (Sn4, Sn15 and Sn79) (yellow clade), *Leptosphaeria maculans* (Lm) and *Zymoseptoria tritici* (Zt) isolates. The branches for race 4 isolates not producing known effectors (Ls13-192, 90-2 and SD20) are highlighted (blue) within the Ptr clade. D) Circular plots show 10 kb regions of absence plotted for the Ptr isolates genomes sequenced using long-read technologies (PacBio and Oxford Nanopore Technology) as compared with the chromosomes of the reference Ptr genome of isolate M4. Isolates are coloured by race. Three regions of interest are highlighted in grey and zoomed at 20x for chromosome 2 and chromosome 1, and 40x for chromosome 9.

Whole genome phylogenetic analysis of Ptr and related ascomycete fungal species clustered into four distinct clades for *Bipolaris* spp., *P. nodorum*, *P. tritici-repentis* and *P. teres* (Fig. 1C). A lower phylogenetic divergence within the individual *Pyrenophora* species (Ptm, Ptr and Ptt) was observed as compared with Bs, Pn and Zt isolates (Supplementary Fig. S8).

To observe regions of absence across the assembled genomes, regions ≥ 10 kb absent for the Ptr isolates were plotted against the reference M4 genome (Fig. 1D). The large horizonal transferred region for ToxA on chr6 was present in all ToxA producing isolates and absent in ToxA non-producing isolates. For the previously reported large Ptr *ToxA* horizontal transfer region, believed to have come from *P. nodorum* (Friesen, Stukenbrock et al. 2006, Manning, Pandelova et al. 2013, Moolhuijzen, See et al. 2018), clear break points on M4 chr6 at the 1,645,874 bp and 1,774,022 bp positions (128 kb insertion) could be determined between isolates producing and not producing ToxA (Fig. 1D and Supplementary Fig. S9). The flanking regions of the breakpoints were highly conserved between the all aligned isolates (Supplementary Fig. S9). A region on chr1 near the 1.47 Mb position was found absent in all non-ToxC producing isolates and the unknown race (ToxC producing) when only looking at long read assemblies (Fig 1D and Supplementary Fig. S10 (plot on left hand side)). The race 4 isolates had more regions of absence, particularly in the distal ends of chromosome 2. A greater number of absent regions were obtained for Illumina sequenced assemblies (Supplementary Fig. S10, plot on right hand side). Regions of variation appear mostly associated with chromosome telomeres and centromeres. In particular, the distal region on M4 chr10 the equivalent of race 5 isolate DW5 chr11 (1,752,563-2,152,826 bp) was mostly unique as compared with races 1, 2, 4 and the unknown race, with fragmented alignments dispersed throughout the last 100 kb of the chromosome surrounding Ptr *ToxB2* (2,152,563-2,152,826 bp) (Supplementary Fig. S11).

### *P. tritici-repentis* mRNA sequence alignment to whole genomes

To ensure a comprehensive search of Ptr genes in the pangenome, predicted mRNA sequences from all isolates were aligned to all the genomes at greater than 90% sequence identity and 90% coverage. The number of alignments and greatest percent identity for each locus were recorded to determine isolate correlations (Fig. 2). Although a closer correlation by gene percent sequence identity could be determined for isolates that were Illumina or PacBio sequenced, a distinct grouping for Alg130, T199 and T205, and a grouping of the European isolates, was evident. Furthermore, the race 4 isolates 90-2 and SD20 were less correlated to all the remaining isolates (Fig 2A). Based on gene counts (copy number) three distinct groups were observed, for long read-sequenced, European Illumina-sequenced and Australian/North-African/North-American Illumina-sequenced isolates (Fig 2B). However, the three race 4 isolates (Ls13-192, 90-2 and SD20) were outliers.

**Fig. 2.**
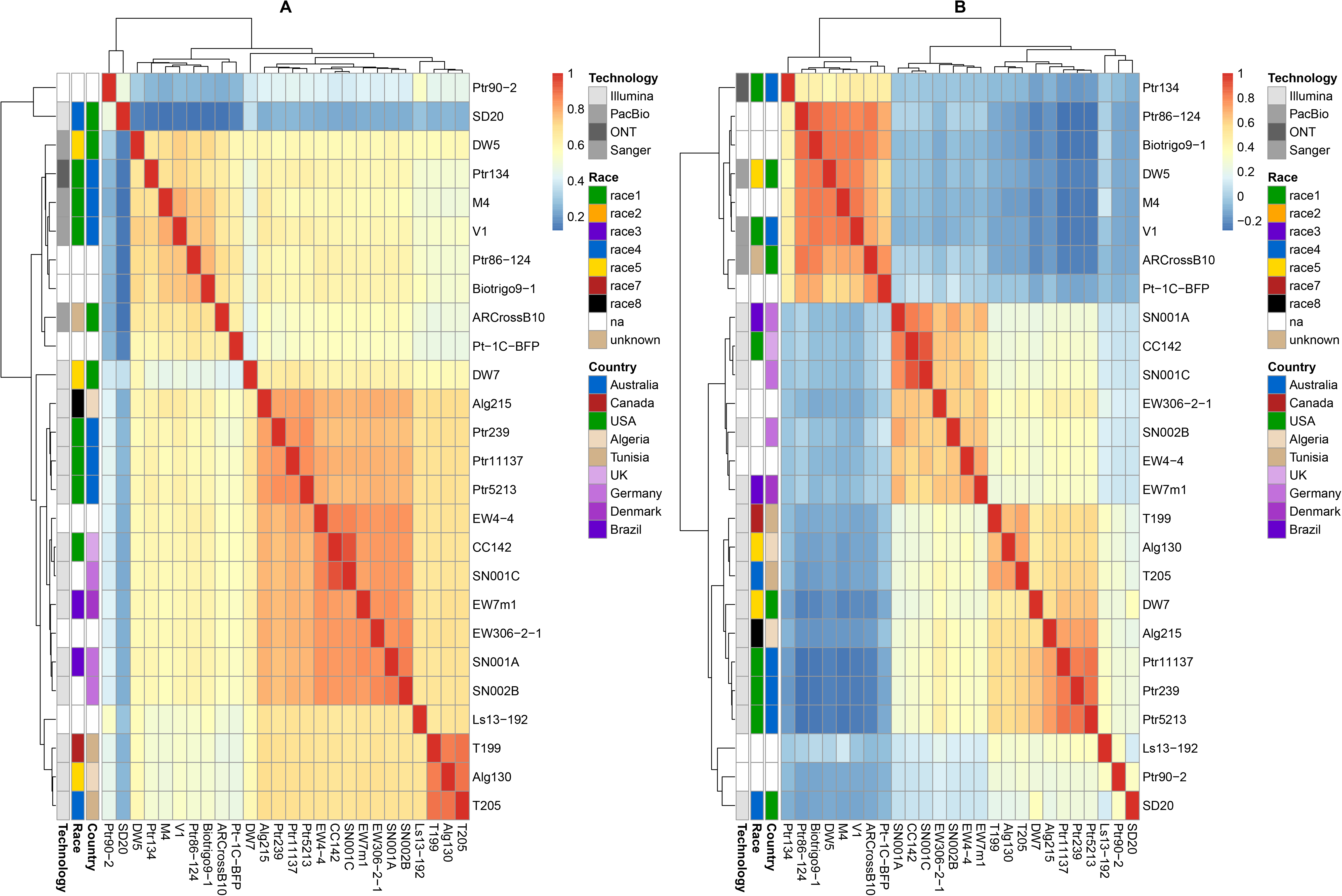
Ptr pangenome predicted mRNA correlation plots for gene sequence percentage identity (A) and gene copy number (B). Ptr isolates from Illumina sequencing (Alg130, T199, T205, Alg215, CC142, EW306-2-1, EW4-4, EW7m1, DW7, Pt-1C-BFP, Ptr239, Ptr11137, Ptr5213, SD20, SN001A, SN001C, SN002B), PacBio Technologies (86-124 (Ptr86-124), 90-2 (Ptr90-2), AR CrossB10, Biotrigo9-1, DW5, Ls13-192, M4 and V1) and Oxford Nanopore Technologies (Ptr134).

All genes were then filtered for presence/absence variation between ToxC-producing isolates [race 1 (Pt-1C-BFP, CC142, EW4_4 and EW306-2-1), race 3 (EW7m1 and SN001A), race unknown (AR CrossB10) and provisional race 8 (Alg215)] and non-ToxC producing isolates [race 2 (86-124 and Biorigo9-1), race 4 (T205, Ls13-192, 90-2 and SD20), race 5 (DW5 and DW7) and race 7 (T199)] to identify genes that may be related to ToxC production. When only PacBio sequenced genomes were queried, a gene cluster of 16 genes from isolate M4 mRNAs 12743 to 12761 (proteins KAF7566087-KAF7566105) positioned on M4 chromosome 9 within 101,367 - 138,426 bp and 14 single loci genes outside of the cluster were found present in the ToxC producing races (races 1 and unknown) and absent in races not producing ToxC (races 2, 4 and 5) (Supplementary Fig. S12). The region was however absent for the race 1 Oxford Nanopore technology (ONT)-sequenced isolate Ptr134 (Fig. 1D). None of the 30 genes found to be specific to ToxC-producing isolates (based on PacBio technology) had an identified signal peptide or appeared to be part of any predicted biosynthetic gene cluster (Table 3 and Supplementary data 3). A search of the pathogen-host interaction database PHI-base (Urban, Cuzick et al. 2017), which provides expertly curated molecular and biological information on genes proven to affect the outcome of pathogen-host interactions, did however identify four proteins with significant alignments to proteins with classified reduced virulence and lethal phenotypes. The following proteins with reduced virulence phenotype were described as being an indoleacetamide hydrolase (iaaH) involved in auxin biosynthesis and plant hormone metabolism (P06618) in *Pseudomonas savastanoi*, a Non-Ribosomal Protein Synthase (NRPS) (A0A024CHY2) in *Pseudomonas cichorii* and an AMP binding protein (E3QPY3) in *Colletotrichum graminicola*. The protein I1RXA5, classified with a lethal phenotype in *F. graminearum*, appears to be a transcription factor (homeobox).

**Table 3.**
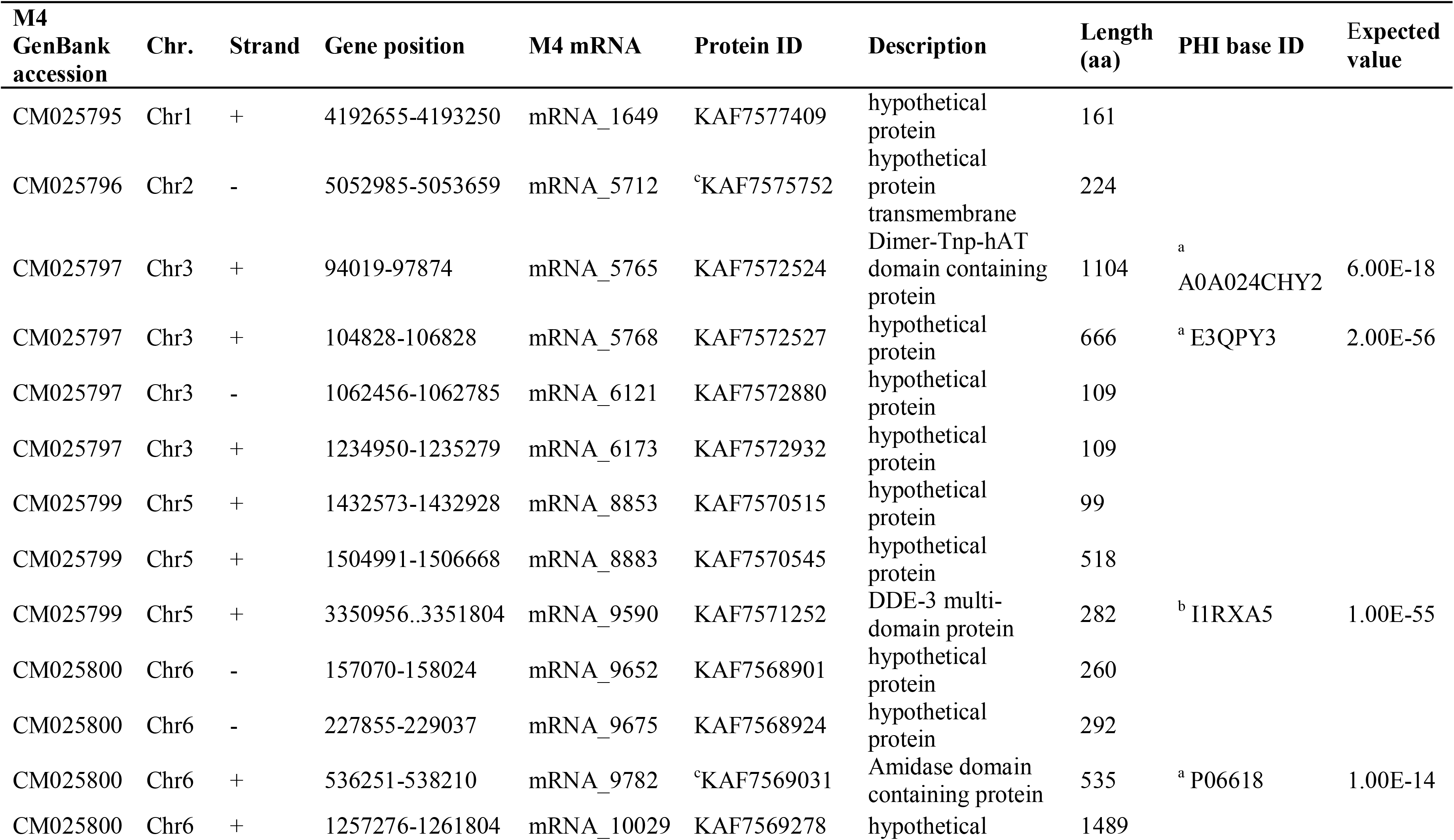
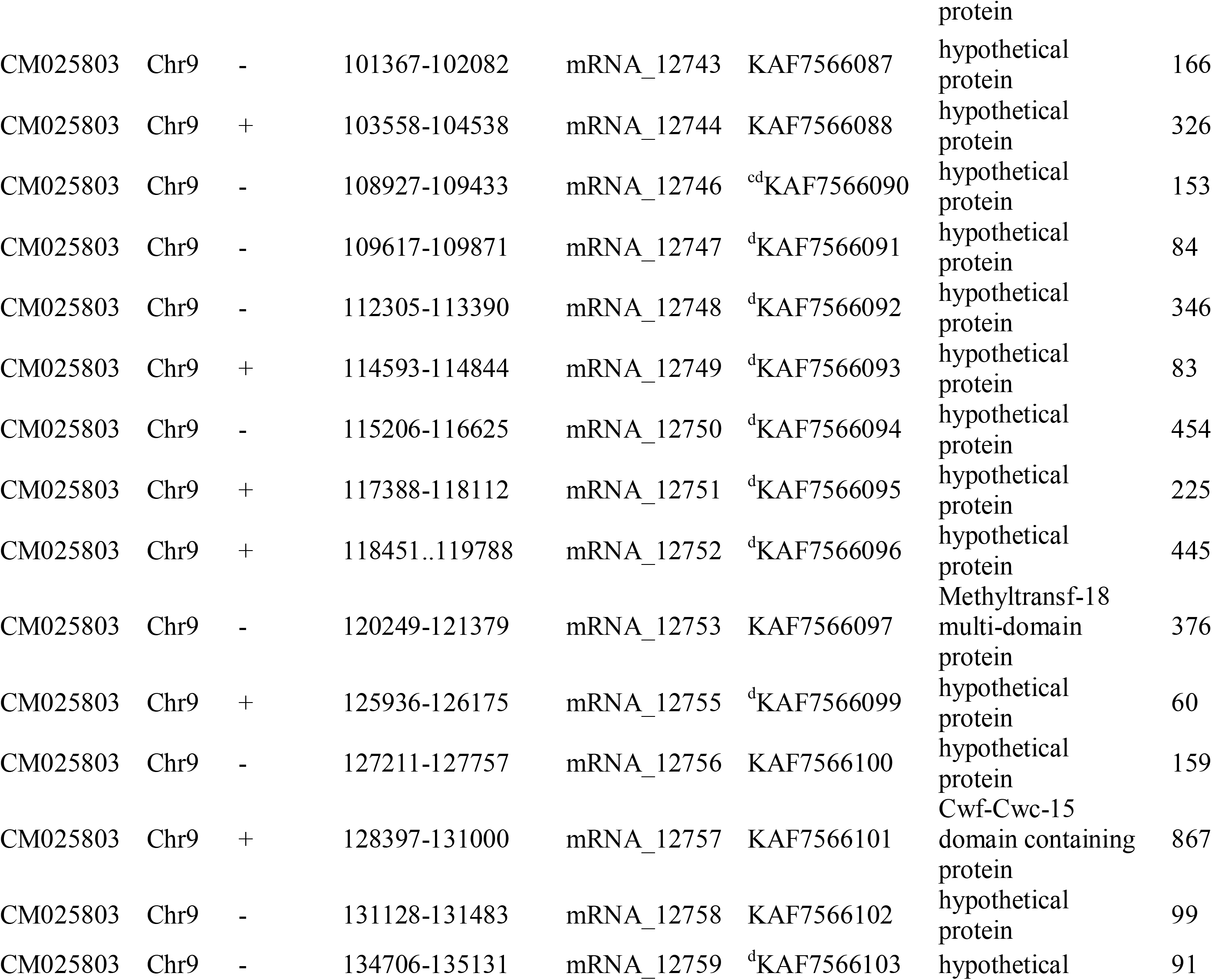
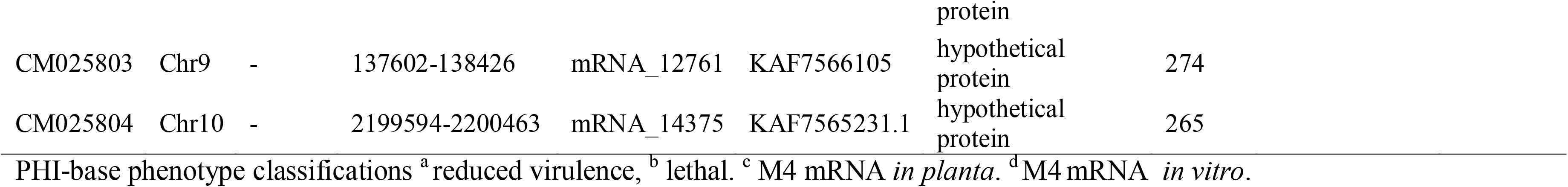
Ptr predicted mRNA sequences identified specific to ToxC-producing isolates (Pacbio sequenced) and PHI-base results.

The genes specific for ToxC-producing isolates (that were PacBio-sequenced) were also searched in previous published *in planta* and *in vitro* RNA-seq data (Moolhuijzen, See et al. 2018) and most of the gene cluster (KAF7566087-KAF7566105) had *in vitro* transcription support (Supplementary Fig. S13). Only the hypothetical transmembrane protein (KAF7575752), amidase containing protein (KAF7569031) and two other hypothetical proteins had *in planta* transcription support during Ptr infection (Table 3).

When all sequenced isolates were considered, only a single locus for a transmembrane protein, an integral membrane component, was identified core to all ToxC-producing isolates, represented by the M4 protein (KAF7575752) on chromosome 2 position 5,052,985-5,053,659 bp (Fig. 1D). This gene was recently identified as *ToxC1*, a gene required but not sufficient for ToxC production in Ptr (Shi, Kariyawasam et al. 2022). A less stringent search for *ToxC1* in all isolates detected the presence of *ToxC1* in the race 2 isolate Biotrigo9-1 genome, which was disrupted by a large insertion of 5,348 bp, positioned at 45,946 to 51,292 bp on contig 12, which disrupted the *ToxC1* protein coding region in the 582-583 bp position. Examination of the 2 kb gene flanking regions of all genomes indicated a further large insertion downstream of the gene in Biotrigo9-1 (Fig. 3). The two large insertions do not have a similar sequence identity, with the insertion downstream of *ToxC1* carrying Gypsy retrotransposon transposable elements (TEs) and the *ToxC1* insertion carrying Copia retrotransposon TEs informed by flanking long terminal repeats (LTRs) (Supplementary Fig. S14).

**Fig. 3.**
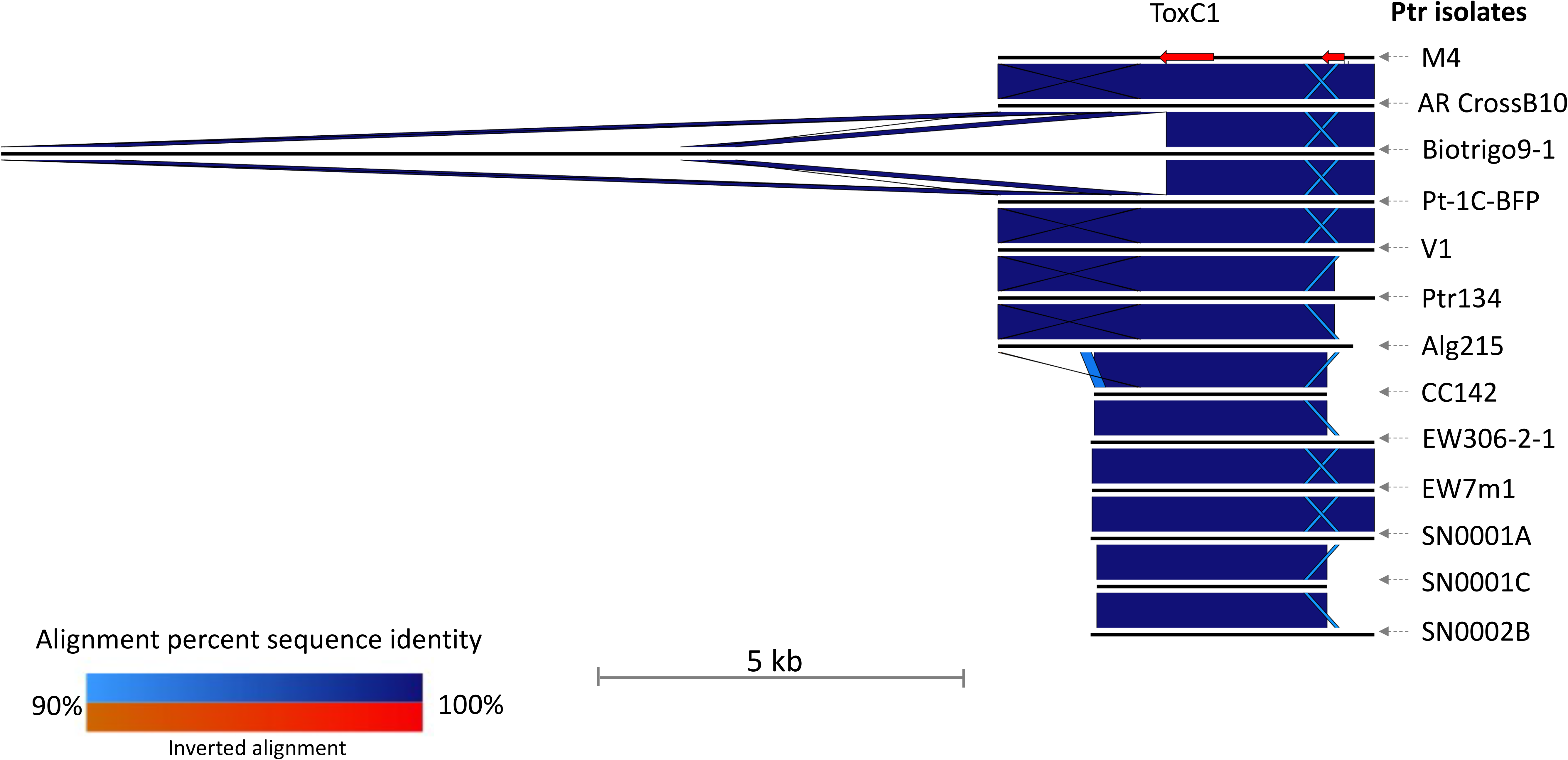
Ptr isolate M4 *ToxC1* locus and 2 kb flanking sequence region alignment to twelve other Ptr ToxC producing isolates. The Biotrigo9-1 *ToxC1* region has two large insertions within and downstream of *ToxC1*. Nucleotide sequence alignments (blue) between the *ToxC1* region for Ptr isolates (top to bottom M4, AR CrossB10, Biotrigo9-1, Pt-1C-BFP, V1, Ptr134, Alg215, CC142, EW306-2-1, EW7m1, SN0001A, SN0001C and SN0002B) (black lines). The M4 genes are shown as red arrows. The light blue alignment segments are regions of low identity among the isolates, while the crossed regions indicate a repeat region in each sequence.

### Analyses of core and ancillary gene sets / protein clusters

To determine core and ancillary protein groups in Ptr, a total of 331,44 predicted protein coding genes from this study and published genomes downloaded from NCBI (see Material and Methods) were clustered. Out of the total number, 328,336 proteins clustered into 14,833 orthologous groups and 3,308 singletons, representing a pangenome for Ptr. A total of 8,503 groups were core (57%) (with all isolates present) and 7,162 orthogroups (48%) consisted entirely of single-copy genes (Supplementary data 4). Overall, for the PacBio sequenced isolates, race 4 isolate 90-2 had the highest percentage of duplicated genes (two copies) (12%) (Supplementary data 4). The percentage of single copy genes for the PacBio sequenced genomes ranged from 76% for M4 to 89% for Ls13-192.

Across the Ptr pangenome (core and ancillary genes), 32,257 (9.6%) genes had a signal peptide of which just over one third (11,911 genes) were predicted to be effectors (EffectorP 3.0 default probability score ≥ 0.5). The EffectorP 3.0 effector probability scores for ToxA and ToxB were 0.702 and 0.93, respectively. The effectors *ToxA* and *ToxB*/*toxb* were identified in orthologous protein groups OG0011421 and OG0011851, respectively.

All predicted effectors protein sequences were then clustered into 738 orthogroups, of which five groups were isolate specific (containing paralogous genes) and 187 were singletons (a single gene). Of the 738 effector orthogroups, only 119 (16%) were core to all isolates and of the core orthogroups 25 orthogroups (21%) had 100% sequence identity. Of the non-core effector groups, 62 orthogroups were absent in the race 4 isolates T205, Ls13-192, 90-2 and SD20.

A comparison of predicted effectors from orthogroups with race 4 absent to those with race 4 present found that the average protein length was shorter (T-test, Wilcoxon adj. *P-value* 2.9e-294) and the effector probability scores were higher (T-test, Wilcoxon adj. *P-value* 1.8-28) (Supplementary Fig. S15).

### Protein tertiary structure analysis of predicted effectors

To identify protein tertiary structure homology, predicted effectors were screened using remote homology detection methods against known protein structures to build 3-D models. Of these, 147 proteins had predicted high confidence tertiary models based on published tertiary protein structures (Phyre2 confidence ≥ 90% and alignment coverage ≥ 90%) (Supplementary data 5). Of the high confidence proteins, a total of 48 and 19 had annotated hydrolase and binding functions, respectively. Five were annotated as effectors, which included Ptr ToxA necrosis effector (KAF7569451) with 100% sequence identity to the Protein Database (PDB) crystal protein structure of ToxA 1ZLD and four elicitor proteins, hrip2 (KAF7578077, KAF7575054, KAF7570798 and KAF757229) based on the crystal structure from *Magnaphorthe oryzae* (PDB 5FID) with sequence identities ranging between 23 - 26%.

The 147 predicted effector proteins with a confident protein tertiary model were then searched against Phi-Base (Urban, Cuzick et al. 2017). A total of 34 proteins had known Phi-Base pathogenicity or reduced virulence hits, of which 11 were plant avirulence determinants, which included ToxA (Supplementary data 5).

To enable the capture of genes that may have been filtered out previously (that may not have a predicted signal peptide), whole genome HMM libraries of M4 were generated for screening using BackPhyre (Kelley, Mezulis et al. 2015). Effector related protein structures were then selected from toxins available in the RCSB PDB for Ptr ToxB (2MM0), toxb (2MM2), ToxA (1ZLD) and SnTox3 (6WES) to identify any other structural homologues and orthologues, respectively. No structural paralogues for ToxA or ToxB were identified in isolate M4 (with confidence levels ≥ 20.0); however, an orthologous structure was identified for SnTox3 with 58% alignment coverage (46 - 138 amino acids) to M4 (protein accession KAF7577476) (104 - 195 amino acids in the alignment) with a confidence score of 95.5 and 34% protein sequence identity (Fig. 4A). This indicated a high confidence that the match between KAF7577476 and the PnTox3 template is a true homology that adopts the overall protein fold and that the core protein is modelled at a high accuracy (2-4 A° from the native, true structure). The 3-D protein structures for SnTox3 (Fig. 4B) and predicted structure for KAF7577476 (Fig. 4C) were then structurally aligned and superimposed with a root mean square distance (RMSD) of 1.14 A° (Fig. 4D).

**Fig. 4.**
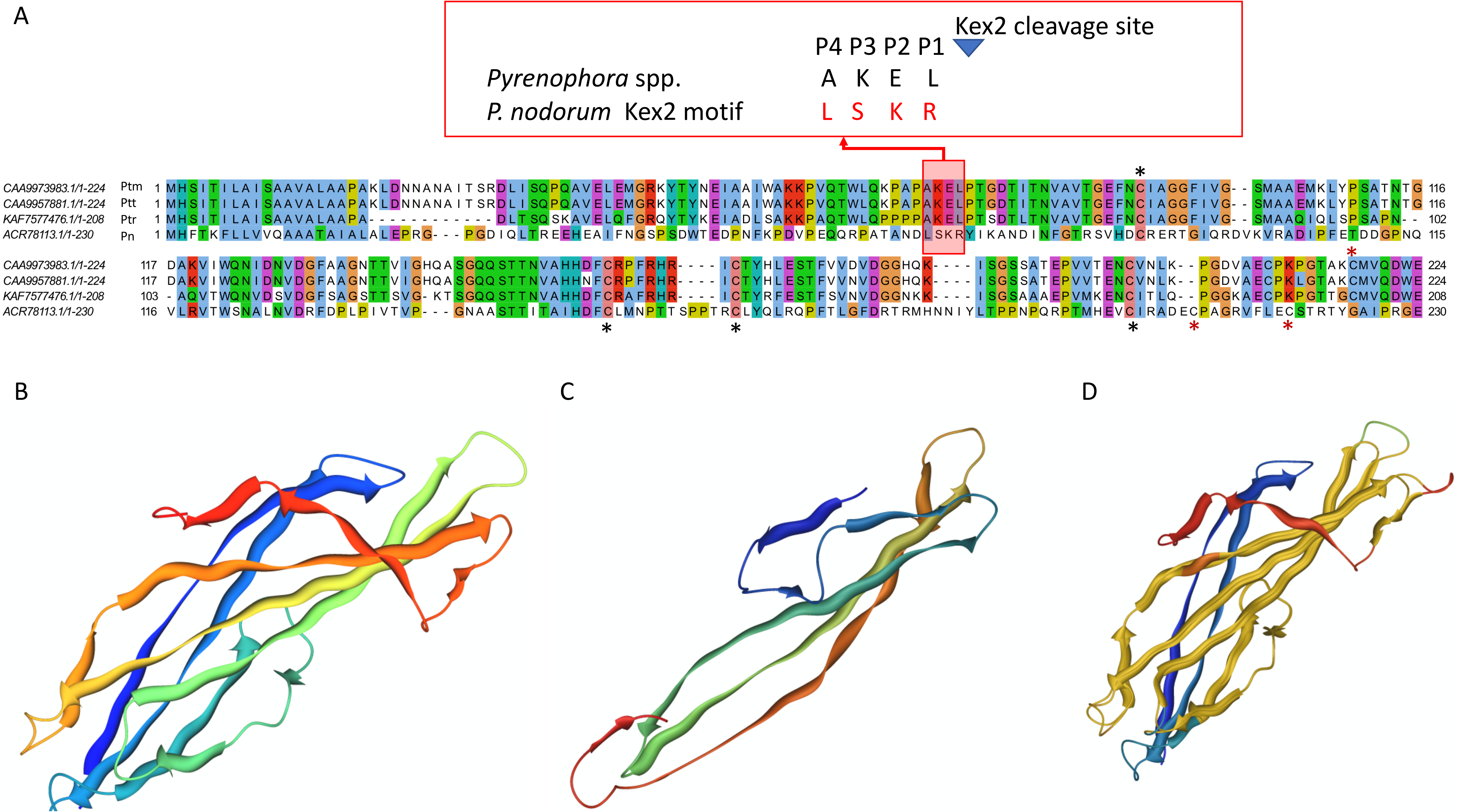
The predicted protein sequence and structural alignments of SnTox3 and the isolate M4 protein KAF7577476. A) Multiple protein sequence alignment of SnTox3, Ptt CAA9973983.1 (W1-1), Ptm CAA9957881.1 (SG1) and Ptr KAF7577476 (M4). The Kex2 motif conservation is shown boxed in red. Only four cysteine residues were conserved across the four species (black asterisks) and those not conserved (red asterisks) are shown below the alignment for *P. nodorum* and above for the *Pyrenophora* spp. B) The known 3-D protein structure for SnTox3 (PDB 6WES). C) The 3-D structure for KAF7577476 as predicted by Phyre2. D) Superimposed structural alignment (yellow) of SnTox3 and KAF7577476 with an RSDM of 1.14 A°.

A TBLASTN sequence search of the M4 isolate predicted protein KAF7577476 against all the Ptr genomes found evidence that the gene encoding this predicted protein is present in all isolates. The KAF7577476 protein sequence was then also searched against the genomes of the related barley necrotrophic fungal pathogens *P. teres* f. *teres* isolate W1-1 (Ptt) and *P. teres* f. *maculata* isolate SG1 (Ptm) (Syme, Martin et al. 2018) and high identity orthologues were also identified: CAA9973983.1 (isolate W1-1) and CAA9957881.1 (isolate SG1), respectively (Fig. 4A). An automated and combinative method for ranking top candidate effector proteins (Predector) (Jones, Rozano et al. 2021) ranked Ptr KAF7577476 in the 262th position, Ptt CAA9973983 as the top candidate (number one) and Ptm CAA9957881 in the 56^th^ position with Predector scores of 1.9, 3.9 and 2.7, respectively (Supplementary data S1). The SnKex2 cleavage motif LSKR (69 - 72 amino acids) of SnTox3 (Outram, Solomon et al. 2021) aligned to AKEL protein residues in the three *Pyrenophora* spp. (Ptr, Ptm and Ptt), where the residue positioned before the cleavage site (P1) is expected to be exclusively an arginine (Arg, R) (Outram, Solomon et al. 2021) (Fig. 4A). Furthermore, the *Pyrenophora* sequences appeared to possess only 4 of the 6 cysteine residues, that form three disulphide bonds (Osbourn 2010), conserved with SnTox3. The predicted apoplastic effector scores for SnTox3, KAF7577476 (Ptr), CAA9973983 (Ptt) and CAA9957881 (Ptm) were 0.573, 0.572, 0.691 and 0.765, respectively.

## Discussion

### *P. tritici-repentis* pangenome analysis

In this study, we present the pangenome of 26 Ptr isolates with a near complete representation of the eight known race categories. Our 15 newly assembled and annotated genomes, along with the 11 previously published genomes, represent a global pangenome of Ptr for major wheat growing regions, with close and distant proximity to the origin of wheat domestication in the Fertile Crescent of western Asia. The repertoire of the known Ptr genes (Moolhuijzen, See et al. 2018) was expanded by 31%, represented by 18,140 non-redundant sequences. This expansion of genes is also observed in other plant fungal species, where a pangenome analysis of 20 *F*. *graminearum* isolates resulted in a 32% gene expansion over the reference isolate (Alouane, Rimbert et al. 2021). The 57% conservation of core orthogroups in Ptr identified here is similar in magnitude to a recent 19 isolate pangenome analysis of the wheat pathogen *Z*. *tritici* (Badet, Oggenfuss et al. 2020), which found 60% of gene orthogroups were core.

A number of Ascomycete genomes, such as Pn, Ptt and Zt, have ‘two speed genomes’, where the genome is compartmentalised into gene-poor AT-rich regions and can have accessory chromosomes. In contrast, Ptr does not appear to have accessory chromosomes and has a GC-equilibrated genome (Dong, Raffaele et al. 2015, Testa, Oliver et al. 2016, Bertazzoni, Williams et al. 2018, Moolhuijzen, See et al. 2018, Syme, Martin et al. 2018). The whole genome phylogenetic analysis clearly showed greater isolate phylogenetic distances within Bs, Pn and Zt isolates as compared with the *Pyrenophora* spp. (Ptm, Ptr and Ptt). However, even with comparatively low phylogenetic distances within Ptr, distinct clades could be detected based on geographic locations. The only exception was isolate Alg215 from Algeria, which clustered with the Australian and American isolates, sharing a large sub-telomeric region in common. This sequence variation, plus a disrupted *ToxB*, set Alg215 apart from the other isolates collected from North Africa. Despite the low whole-genome phylogenetic distances in Ptr, a lower percentage of core orthogroups (57%) was found compared with a previous analyses of 11 isolates (PacBio and Illumina sequenced), which indicated 69% core orthologous groups (Moolhuijzen, See et al. 2018). This suggests that not only has the pangenome complexity risen with an increase in the numbers of isolates sequenced (as expected), but that an increased divergence in Ptr conserved protein domains is apparent.

Although in this analysis only a single gene was identified as specific to all ToxC producing isolates (*ToxC1*), PacBio sequencing identified a potential gene cluster of interest which would be near impossible to identify in Illumina-sequenced genomes, due to the repetitive nature of the region. Interestingly, our analysis found no putative effectors that were core to all isolates, again indicating a large variability within Ptr for this type of gene. Recently, *ToxC1* was functionally validated using a gene knockout approach (Shi, Kariyawasam et al. 2022), where it was found to be required, but not sufficient, for ToxC production. In our study, no clear gene cluster for a secondary metabolite or Ribosomally synthesized and post-translationally modified peptides (RiPPs) was identified, in part due to the positioning of the *ToxC* locus within the complex subtelomeric region of chromosome 2 (Shi, Kariyawasam et al. 2022), which despite long read sequencing still remains a problematic region to resolve. The presence of *ToxC1* in a non-ToxC producing isolate (Biotrigo9-1) was surprising, and raises more questions regarding the evolution and/or origin of ToxC production. It is possible that the large *ToxC1* insertion by a LTR retrotransposon has disrupted the production of ToxC in Biotrigo9-1 and that those remaining gene(s) involved in ToxC production are present.

The divergence of Ptr race 4 isolates (that do not produce the known effectors on wheat) from isolates that produce known effectors was clearly shown, except for T205. The genome sizes and gene duplication rates of the two race 4 isolates (Ls13-192 and 90-2) also revealed a complexity that was unexpected. Race 4 was first described 30 years ago (Lamari and Bernier 1989, Lamari and Bernier 1991) as a nec^-^ chl^-^ pathotype (avirulent) on the set of differential wheat lines, and has since been reported but not as frequently as the other races from collections of Ptr across different wheat growing regions. A recent study by Guo et al. (2020) showed that despite the inability of race 4 isolates to induce tan spot symptoms on the differential wheat lines, four race 4 isolates (Ls13-14, Ls13-86, Ls13-192 and Ls13-198) from North Dakota in the USA induced varying degrees of disease reactions upon inoculation on tetraploid (durum) wheats (Guo, Shi et al. 2020). This may well provide an explanation for the observed distinction of Ls13-192 from the other race 4 isolates (90-2 and SD20) in the whole genome phylogenetic clustering and gene correlation analyses, since unlike Ls13-192, SD20 and 90-2 have not been reported to be virulent on durum wheat. Furthermore, as *ToxA* and *ToxB* were absent, race 4 isolate T205 is unlike the new virulence type that lacked ToxA and ToxB gene expression on bread wheat differentials but produced necrosis in durum wheat (Benslimane 2018).

While it was unexpected that a race 4 isolate (90-2) had the highest percentage of genes predicted as effectors, this appeared to be the result of a genome-wide expansion of gene copies (which included predicted effectors). It is possible that although the predicted effectors in race 4 isolates may have not have a pathogenic role in bread wheat, they may play a role in another system.

We report, for the first time, an identical *toxb* (non-toxic homologue of *ToxB*) copy in a race 4 genome (2 genes in 90-2). As each *toxb* are on separate contigs, it is not possible to identify if they are co-located. We can, however, speculate that based on the difficulty in assembling the *toxb* regions, they may lie in a subtelomeric chromosome location similar to the multicopy *ToxB*, which was shown to be nested in the complex subtelomeric chromosomal regions of the DW5 genome (Moolhuijzen, See et al. 2020). It is increasingly believed that effector genes are located in transposon-rich and gene-sparse subtelomeric regions of the pathogen genome, allowing opportunity for gene duplication events and thereby contributing to the evolution of virulence diversity. We also show no conservation between the different races for the *ToxB* locus or flanking regions. The sequence variation in the chromosomal centromeric and telomeric regions shown in our whole genome alignments, indicates that these regions are indeed hot spots for diversity. It is furthermore interesting that one of the North Africa isolates (Alg 215) had a truncated *ToxB* gene with a nonsynonymous mutation within the coding region, which may have resulted in a weak chlorosis phenotype on the wheat differential lines. We believe that this is the first report of a *ToxB* nonsynonymous mutation in Ptr. The large *ToxA* horizontal transfer region previously identified (Manning, Pandelova et al. 2013, Moolhuijzen, See et al. 2018) was shown to be absent in all non-ToxA producing isolates and clear insertion breakpoints were identified in all ToxA producing isolates.

In this study, a pangenome approach was undertaken to approximate the complete gene repertoire of the species to capture all gene variations (percentage identity and copy number) and identify candidate genes specific for ToxC-producing races.

### A new *Pyrenophora* resource to identify protein structural homologues

Pathogenic fungi possess large effector repertoires that are dominated by hundreds of small secreted proteins only related by protein tertiary structures (3-D structure) (de Guillen, Ortiz-Vallejo et al. 2015). The prediction of new effector candidates that are not the result of horizontal gene transfer is therefore complicated.

To conduct a comprehensive whole genome search of protein tertiary structures an *in silico* screening was employed using BackPhyre (Kelley, Mezulis et al. 2015). We present here the first necrotrophic fungal pathogen publicly available through BackPhyre (Kelley, Mezulis et al. 2015) for effector and other protein tertiary structure searches, providing further annotation evidence for a number of hypothetical genes. In this pangenome screen of proteins, no other ToxA or ToxB-like paralogues were identified based on structural similarity in Ptr.

Overall, the use of protein three-dimensional structure modelling improved the identification of a number of proteins which included effector candidates potentially involved in pathogenicity.

### *In silico* protein structural analysis reveals a natural homologue to *SnTox3* in *Pyrenophora*

We report here for the first time a distant *SnTox3* natural homologue in *Pyrenophora*. We showed conserved structural homology between SnTox3 and *Pyrenophora* proteins that lacked conservation in the R residue position of the Kex2 motif (LXXR) and the full set of cysteine residues forming the three disulphide bonds in SnTox3. SnTox3 is a pro-domain containing effector, where the signal peptide and pro-domain are removed (cleaved by the Kex2 protease) to produce a more potent protein that activates host cell death (Snn3) (Outram, Sung et al. 2021). The Kex2 cleavage motif (LXXR) has the following residue preferences, a Leucine (L, Leu) or any other aliphatic residue, any residue X as it does not interact with Kex2, Lysine (K, Lys) but has other possible residues Lys > Arginine (R, Arg) >Threonine (T, Thr) > Proline (P, Pro) > Glutamic acid (E, Glu) > Isoleucine (I, Ile) (X) and exclusively an arginine (R, Arg) before the cleavage site (Outram, Solomon et al. 2021). Here we found the conserved *Pyrenophora* motif (AKEL) did not conformed to the Kex2 cleavage motif (LXXR) in two residue positions that included the exclusive arginine residue.

Interestingly, the Ptr structural homologue to SnTox3 was in all isolate races, unlike the non-active *toxb* that only occurs in non-pathogenic race 4 isolates (not producing known effectors). As no *in planta* gene expression for the Ptr homologue of *SnTox3* was detected and the protein sequence had a low effector prediction ranking, we believe it may not be an effector candidate in the wheat-pathogen system. However, conversely in Ptt, as the structural homologue to *SnTox3* is expressed during barley infection (Moolhuijzen, Lawrence et al. 2021) and was ranked as the top candidate effector, we believe further investigation is warranted. Here, we propose that the identification of a SnTox3 structural homologue in *Pyrenophora* (Ptm, Ptr and Ptt) could be part of a structurally defined family and are phylogenetically related to *SnTox3*, as observed for the *<u>M</u>. oryzae* Avirulence (<u>A</u>vrs) and To<u>x</u>B (MAX-effector proteins) (de Guillen, Ortiz-Vallejo et al. 2015).

In conclusion, the new genomic resources presented here improve the pangenome representation of Ptr and provide putative effector candidates based on structural modelling and ranking specific to effector producing isolates. These resources can be used to monitor Ptr variations potentially involved in pathogenicity. As Ptr is commonly shown to infect wheat in combination with other necrotrophic pathogens (Justesen, Corsi et al. 2021), the future ability to simultaneously monitor such changes in multiple necrotrophic species may enhance pathogen monitoring activities within a wider framework of crop protection activities.

## Funding

This work was generously supported through co-investment by the Grains Research and Development Corporation (GRDC) and Curtin University (project code CUR00023), as well as the Australian Government National Collaborative Research Infrastructure Strategy and Education Investment Fund Super Science Initiative. This project was also supported by the Agriculture and Food Research Initiative competitive grants program (award number 2016-67014-24806) and the National Institute of Food and Agriculture, United States Department of Agriculture (USDA) Hatch project (ND02234) to ZL. JC, JT and LJ were supported by the ‘Efectawheat’ project funded within the framework of the 2^nd^ call ERA-NET for Coordinating Plant Sciences by British Biological Sciences Research Council (BBSRC) grant BB/N00518X/1 to JC and the Danish Council of Strategic Research grant case number 5147-00002B to LJ. The funders had no role in the design of the study; in the collection, analyses, or interpretation of data; in the writing of the manuscript, or in the decision to publish the results.

## Acknowledgements

We thank the Australian grain growers for their continued support of research through the Grains Research and Development Corporation (GRDC) and the Australian Government National Collaborative Research Infrastructure Strategy (NCRIS) for providing access to Pawsey Supercomputing under a National Computational Merit Allocation Scheme (NCMAS), Nectar Research and Pawsey Nimbus Cloud resources. We would also like to acknowledge Professor Richard Oliver who was key in the setting up the collaborators in this study as part of the EfectaWheat project.

## Competing interests

The authors declare no competing interests.

## Author contributions

Conceptualisation ZL and CSM; methodology, PM, PTS and HP; formal analysis, PM, PTS, GS and HP; investigation, PM; project resources JC, JT, SS, HB and LJ; writing - original draft preparation, PM; writing – review and editing, PM, PTS, ZL, GS, HB, JC, LJ, SS and CM. All authors have read and agreed to the published version of the manuscript.

## Data availability

All data generated or analyzed during this study are included and can be accessed in this published article (and its supplementary files). The sequence data has been deposited in the DDBJ/ENA/GenBank under accession numbers JAAFOX000000000, JAHCSW000000000, JAHCYZ000000000, NRDI02000000 (version 2), PSOO00000000-PSOU00000000 and RXHK00000000-RXHN00000000.

## Supporting Information

### Supplementary data

Supplementary data 1 XLSX. Predicted effector genes for *Pyrenophora tritici-repentis* genomes.

Supplementary data 2 PDF. List of URLs for publicly available isolate genomes downloaded from NCBI for this study.

Supplementary data 3 XLSX. Predicted biosynthetic gene clusters for *Pyrenophora tritici-repentis* genomes.

Supplementary data 4 XLSX. Orthologous protein clusters for *Pyrenophora tritici-repentis* genomes.

Supplementary data 5 XLSX. Phyre2 three-dimensional protein modelling for *Pyrenophora tritici-repentis* predicted effector proteins.

### Supplementary figures

Supplementary Fig. S1. BUSCO quantitative assessment of the completeness of genome assemblies in terms of expected gene content.

Supplementary Fig. S2. Ptr *ToxB* and *toxb* nucleotide sequence alignments.

Supplementary Fig. S3. Ptr ToxB and toxb protein sequence alignments.

Supplementary Fig. S4. Ptr Alg215 isolate partial *ToxB* sequence alignments show Alg215 *ToxB* is truncated at the 5’ end of the sequence. A) The first 162 nucleotide bases of Alg215 scaffold 03337 sequence aligned to the *ToxB* coding sequence (CDS) (99-261 bp). A *ToxB* single nucleotide polymorphism (SNP; at the 149 bp position) shows a thiamine nucleotide change to guanine (T > G). B) The Alg215 *ToxB* region protein translated (1-94 aa) aligned to the ToxB protein sequence (1-87 bp). A nonsynonymous amino acid residue change (I>R) at ToxB residue position 50 is beyond the ToxB signal peptide cleavage site between amino acid positions 23 and 24.

Supplementary Fig. S5. Ptr plant leaf infection assays to identify isolate ToxA production by the development of necrosis symptoms on the differential wheat cultivar Glenlea (left hand side) and ToxB production by chlorosis symptoms on the differential wheat line 6B662 (sensitive) (right hand side).

Supplementary Fig. S6. Ptr plant infection assays to confirm no symptoms on response to the differential wheat line Auburn (insensitive) (left hand side) and ToxC production by chlorosis symptoms on the differential wheat line 6B365 (right hand side).

Supplementary Fig. S7. Closer examination a large 1 Mb distal region on isolate M4 contig 1 and many smaller regions were absent in isolates Alg130, T199 and T205 but present in Alg215.

Supplementary Fig. S8. Phylogenetic analysis of publicly available ascomycete genomes downloaded from NCBI.

Supplementary Fig. S9. Ptr isolate M4 ToxA horizontal transfer and flanking genomic region (400 kb) alignments for race 1, 2, 4, 5 and unknown races. Break points are displayed for the large 128 kb insertion in the M4 isolate. The *ToxA* horizontal transfer region is absent in all non-ToxA producing races (race 4, 5 and unknown) and present in ToxA producing races (race 1 and 2).

Supplementary Fig. S10. Circular plots show 10 kb regions of absence for Ptr isolates as compared to M4, coloured by race. The left plot shows assembled isolate genomes sequenced from long-read technologies, PacBio and Oxford Nanopore Technology. The right plot displays all the genomes.

Supplementary Fig. S11. DW5 chromosome 11 *ToxB2* and flanking genomic region alignments for race 1, 2, 4, 5 and unknown races. Slide 1, EasyFig Blastn alignments for 400 kb region. Top to bottom DW5 aligned to race 4 (90-2 and Ls13-192), race unknown (AR CrossB10), race 2 (86-124 and Biotrigo9-1) and race 1 (V1, M4 and Ptr134). Slide 2, NUCmer sequence dot plot for 200 kb region.

Supplementary Fig. S12. Ptr loci found only in PacBio sequenced ToxC-producing isolates. Loci are absent in non-ToxC producing Ptr isolates Ls13-192 (race 4), 86-124 and Biotrigo9-1 (race 2), DW5 (race 5) and are present in ToxC producing isolates AR CrossB10 (AR, race unknown), V1 and M4 (race 1) isolates. M4 isolate had multiple gene copies, while some genes were absent in Pt-1C-BFP (race 1).

Supplementary Fig. S13. RNA expression for *Pyrenophora tritici-repentis* isolate M4 for the M4 ToxC producing isolate specific gene cluster 64 kb region (red). The read coverage and alignments show RNA expression *in vitro* (top) and *in planta* (below) on chromosome 9 (CM025803.1).

Supplementary Fig. S14. Ptr isolate Biotrigo9-1 *ToxC1* genomic region on contig 12 (43,362-60,409 bp) shows *ToxC1* is disrupted by a large single insertion (5,348 bp in size) positioned from 45,946 to 51,292 bp that carries a nested long terminal repeat (LTR) retrotransposon and the motifs for the Copia retrotransposon transposable element (TE).

Supplementary Fig. S15. Ptr pangenome predicted effectors specific to race 4 and specific to known effector producing isolates (non-race 4). A) Boxplot of protein lengths. B) Boxplot of effector probability scores. C) Violin plot of lengths. D) Violin plot of effector probability scores.

## Notes

### Competing Interest Statement

The authors have declared no competing interest.

### Summary of Updates

Minor edits

