## Supplementary figures and images for "A global pangenome for the wheat fungal pathogen *Pyrenophora tritici-repentis* and prediction of effector protein structural homology"

### Supplementary Fig. S1

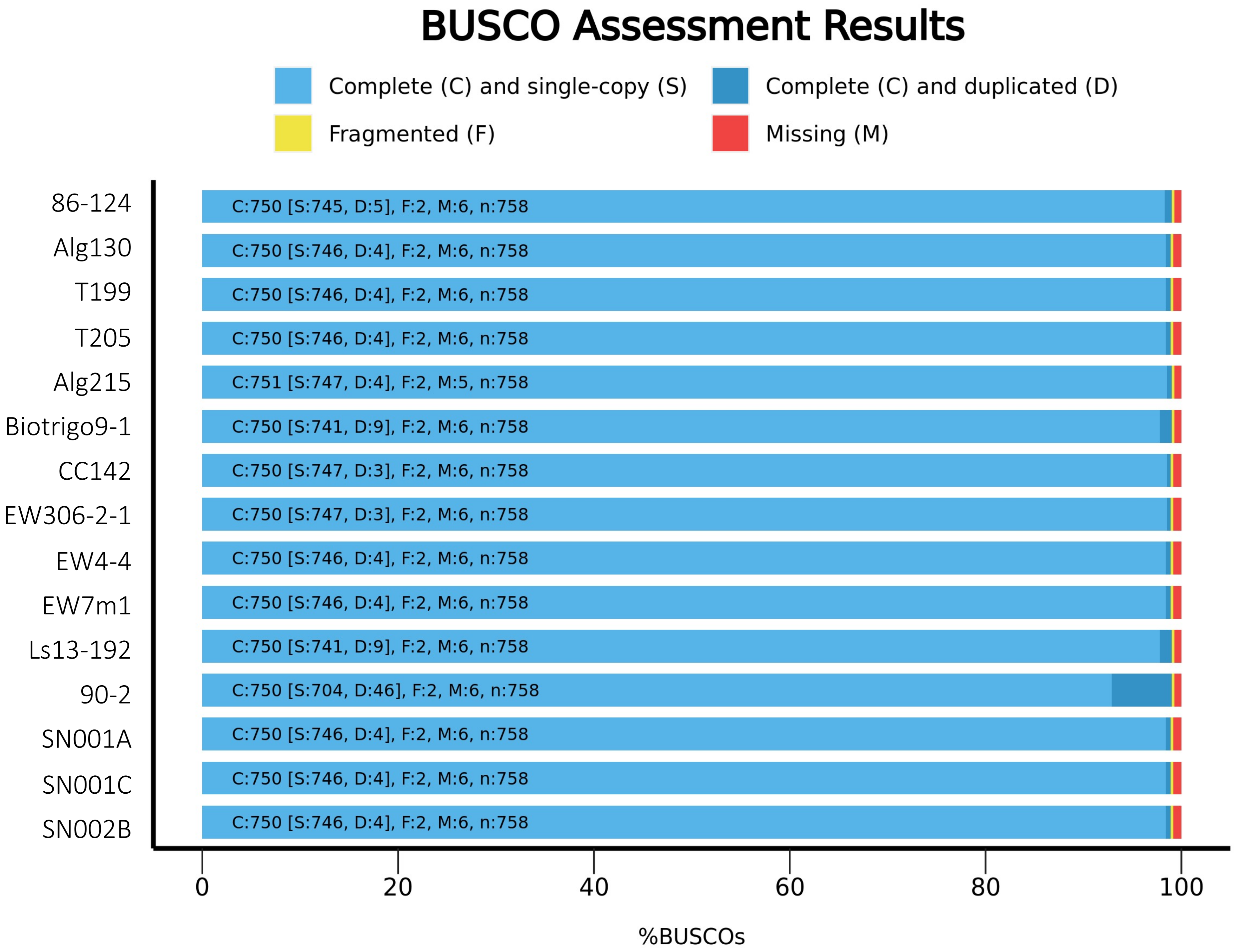

### Supplementary Fig. S2

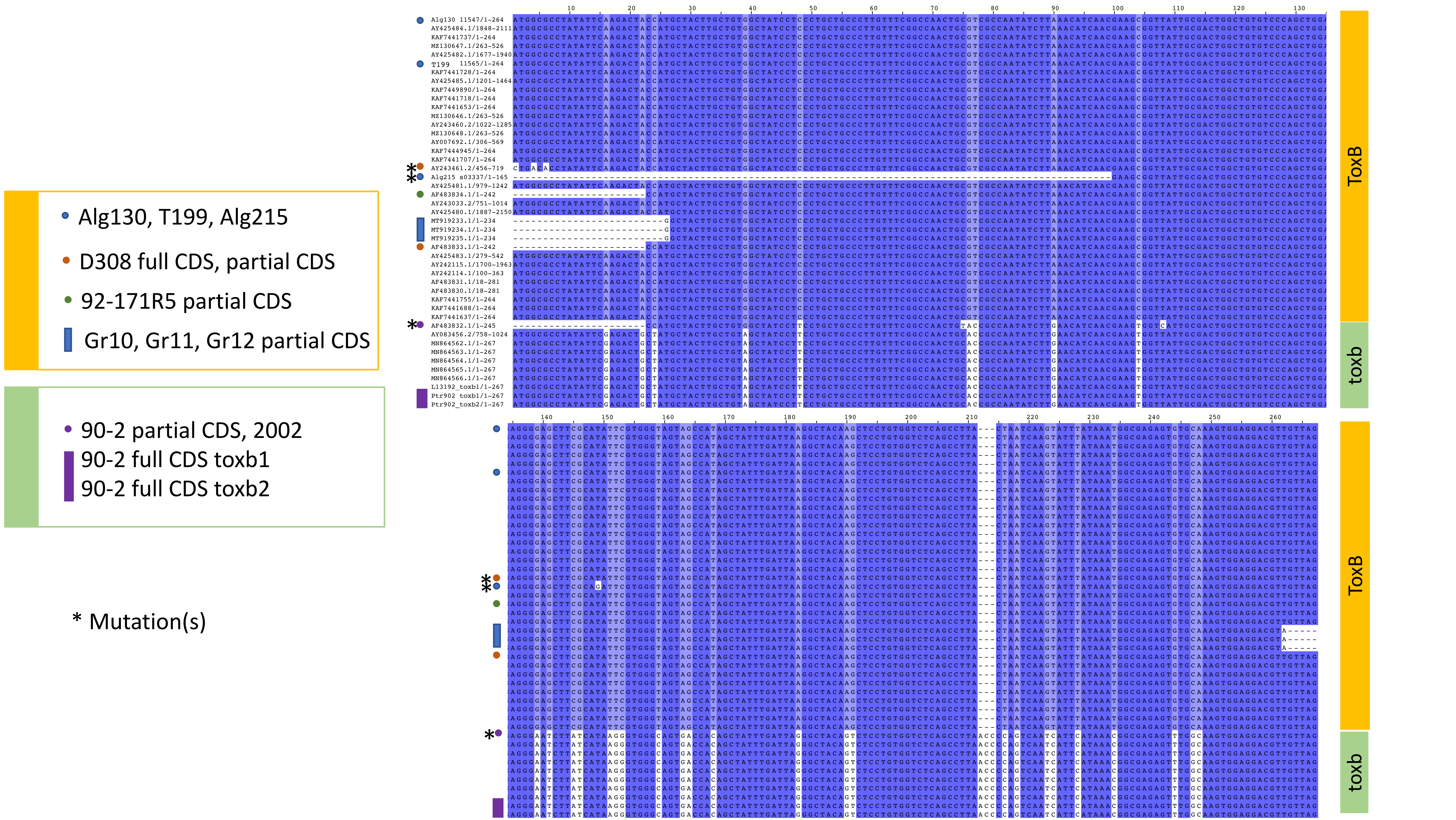

### Supplementary Fig. S3

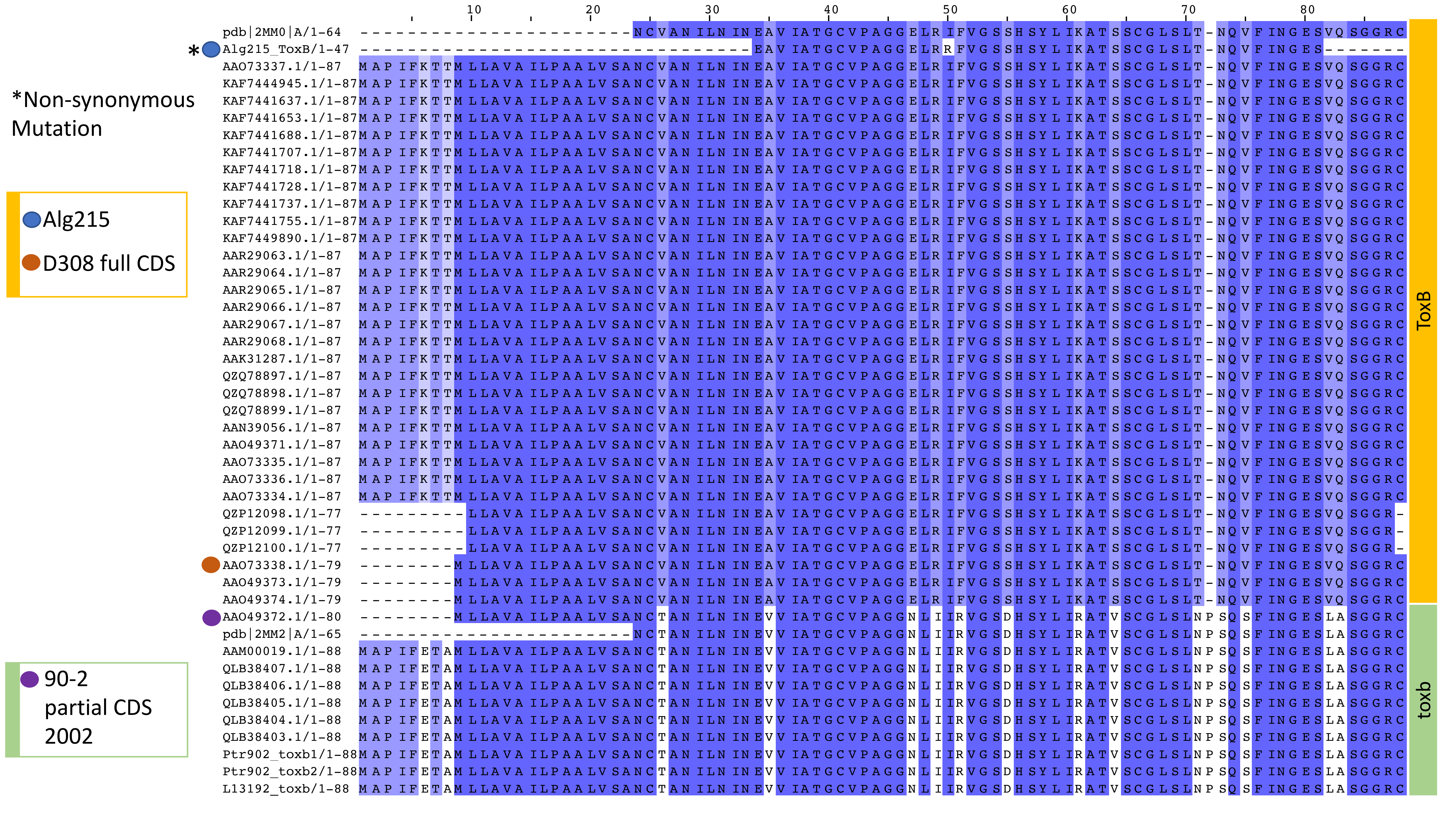

### Supplementary Fig. S4

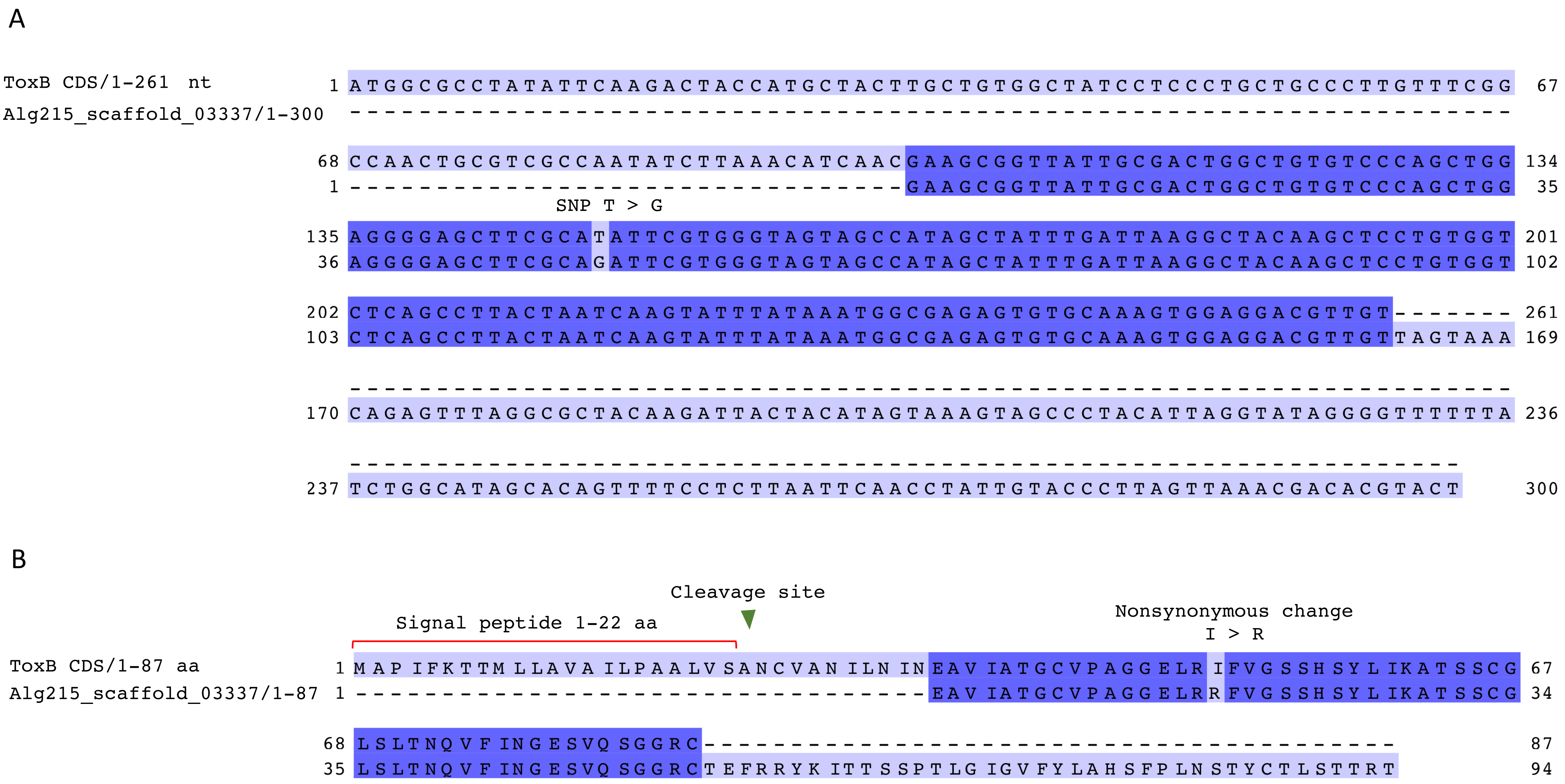

### Supplementary Fig. S5

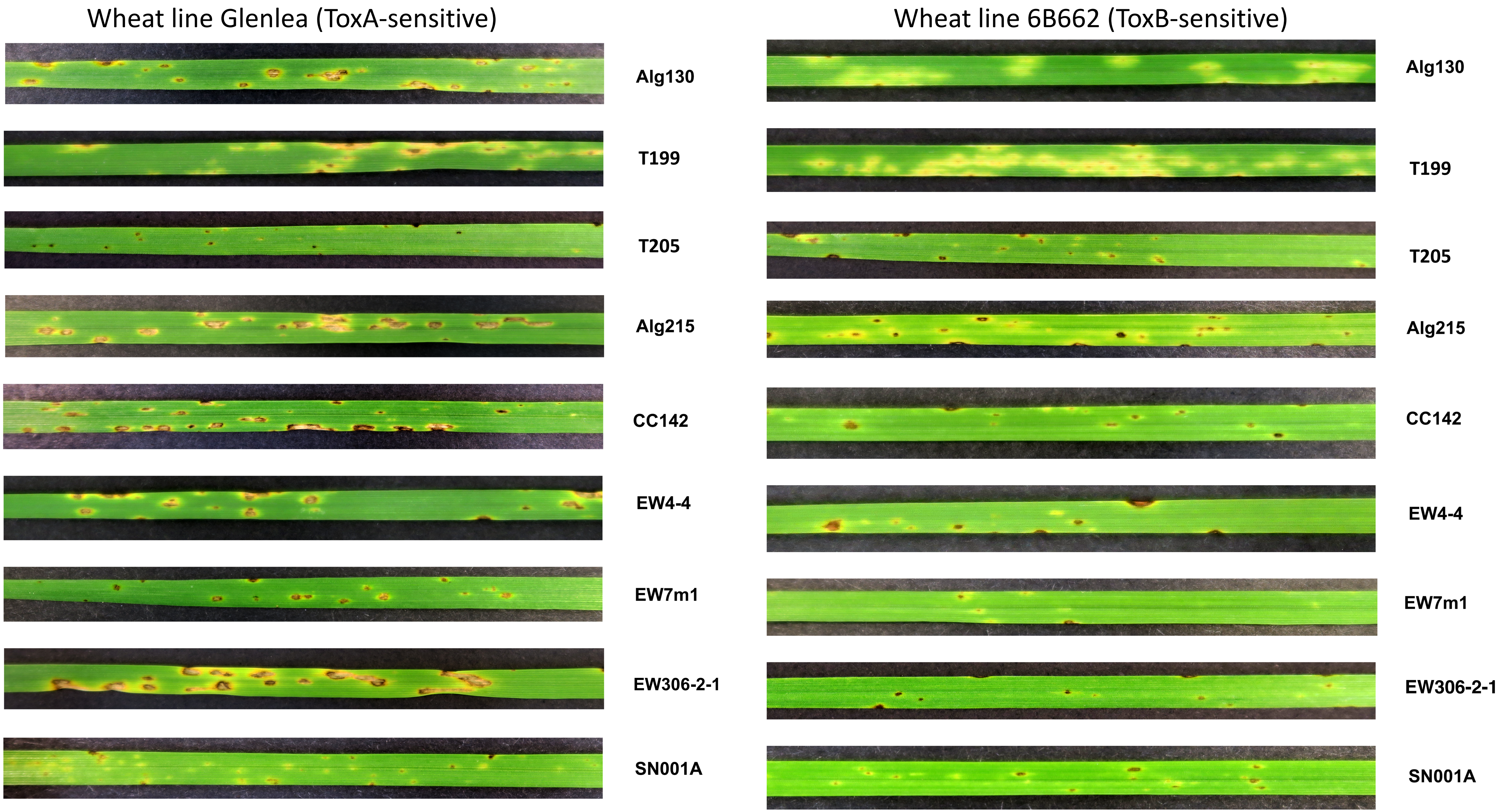

### Supplementary Fig. S6

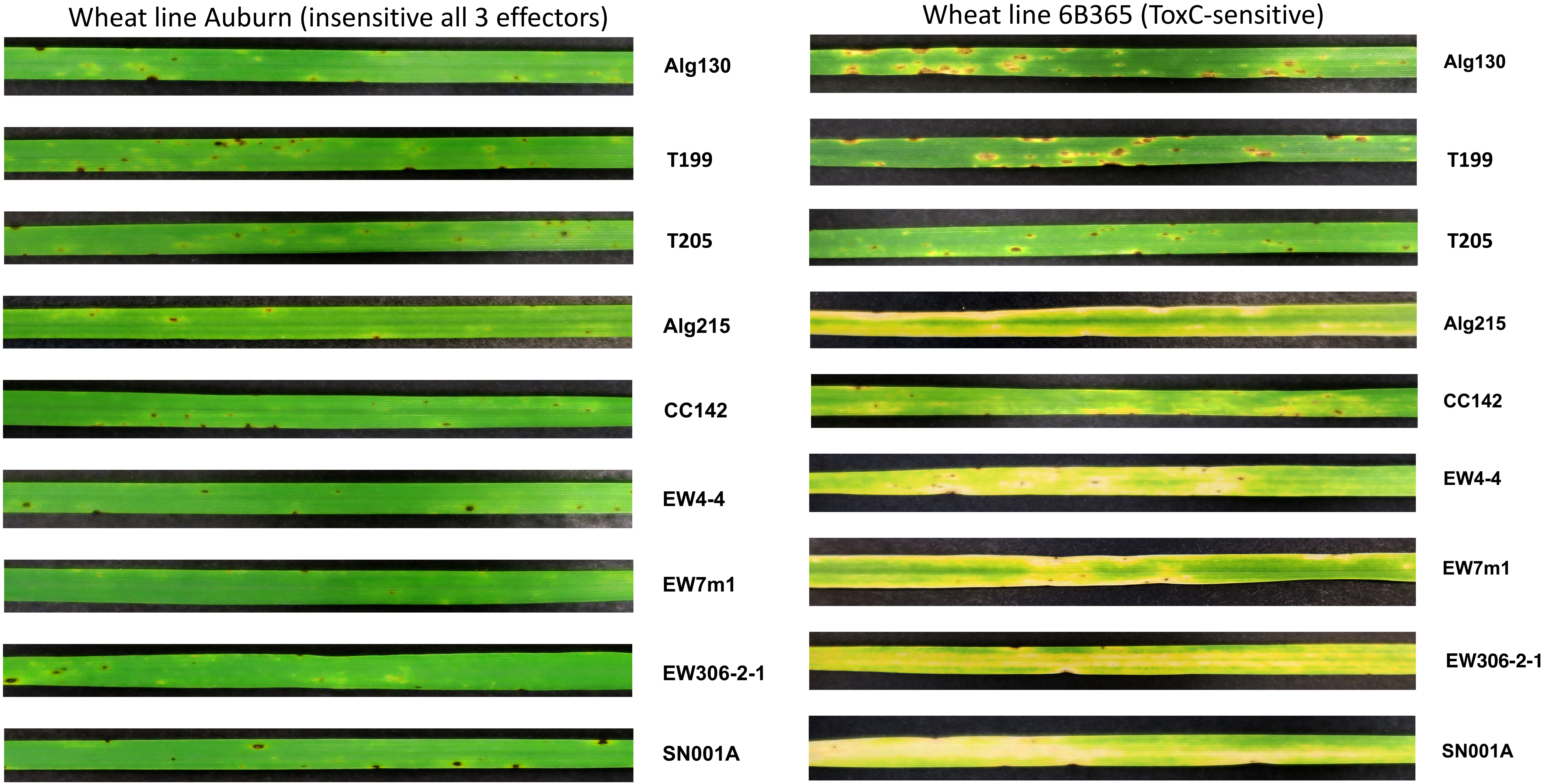

### Supplementary Fig. S7

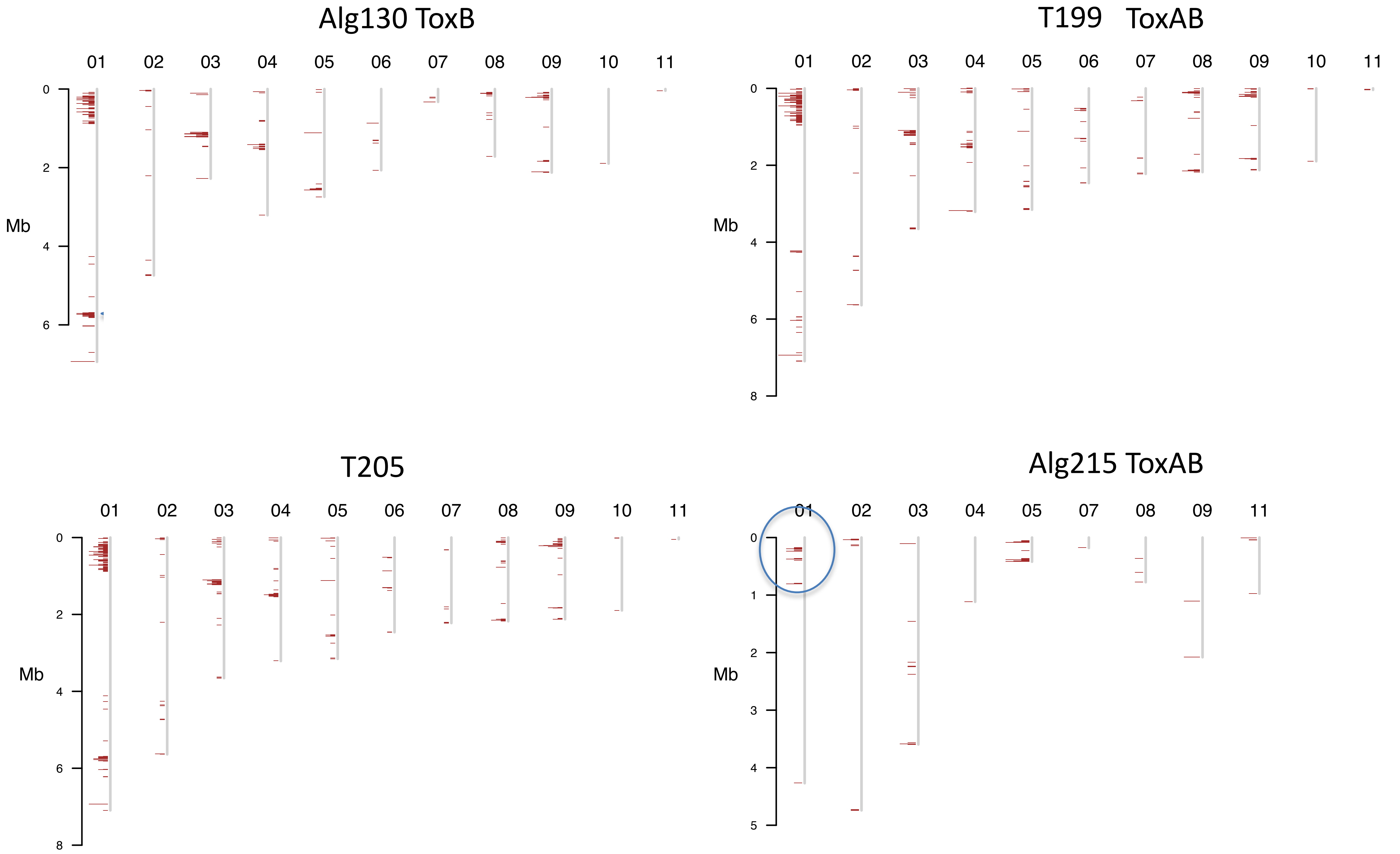

### Supplementary Fig. S8

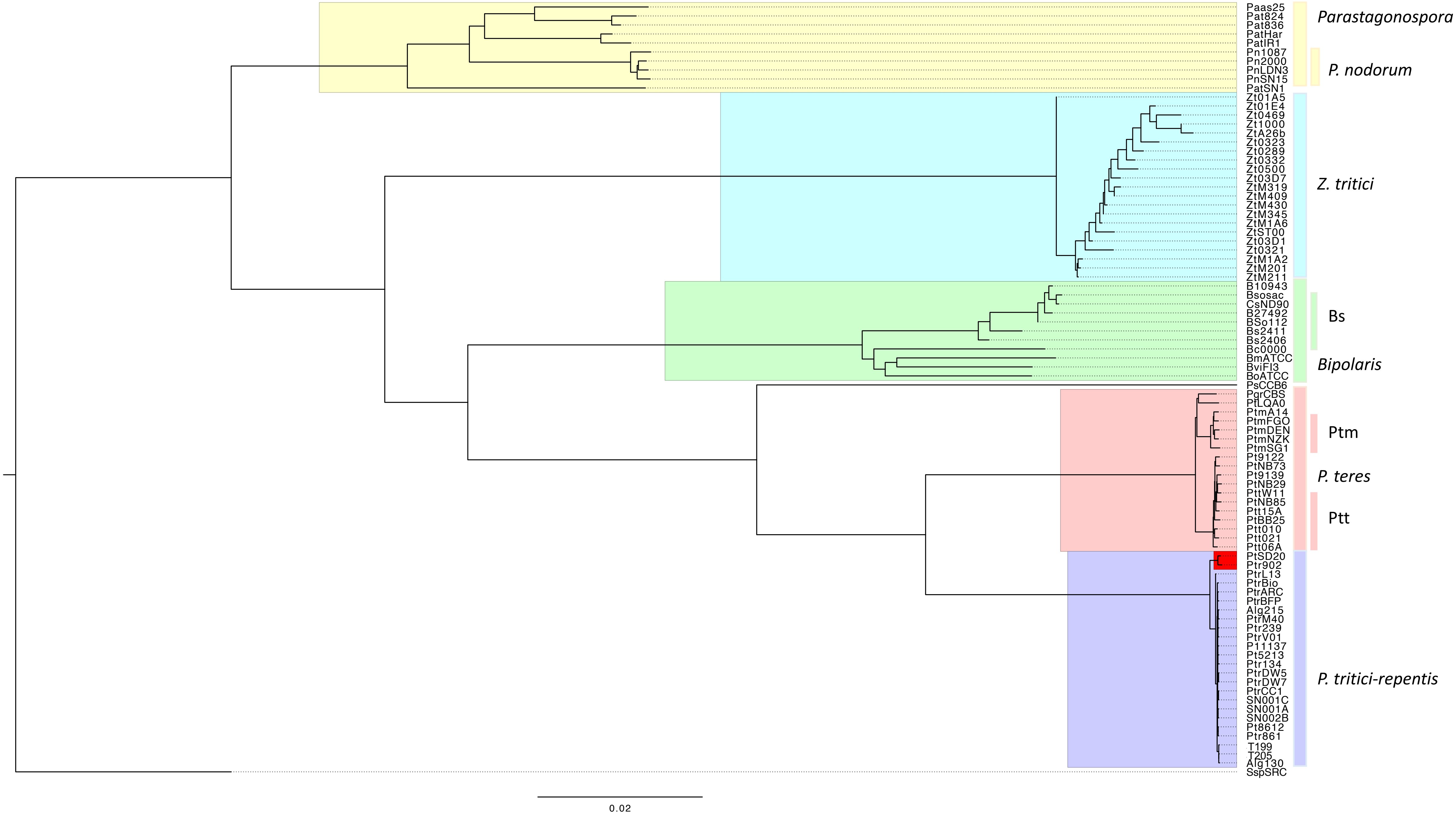

### Supplementary Fig. S9

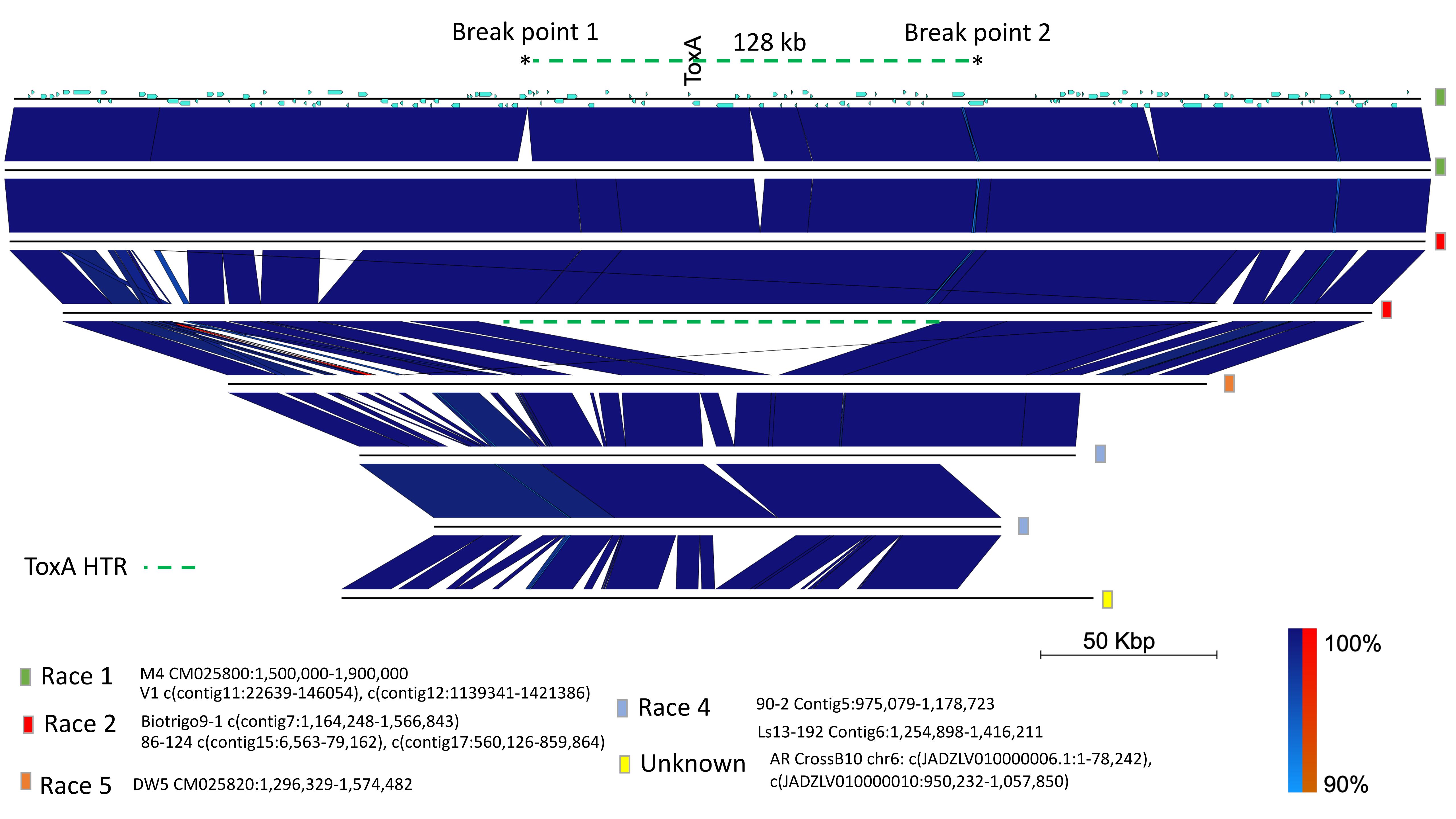

### Supplementary Fig. S10

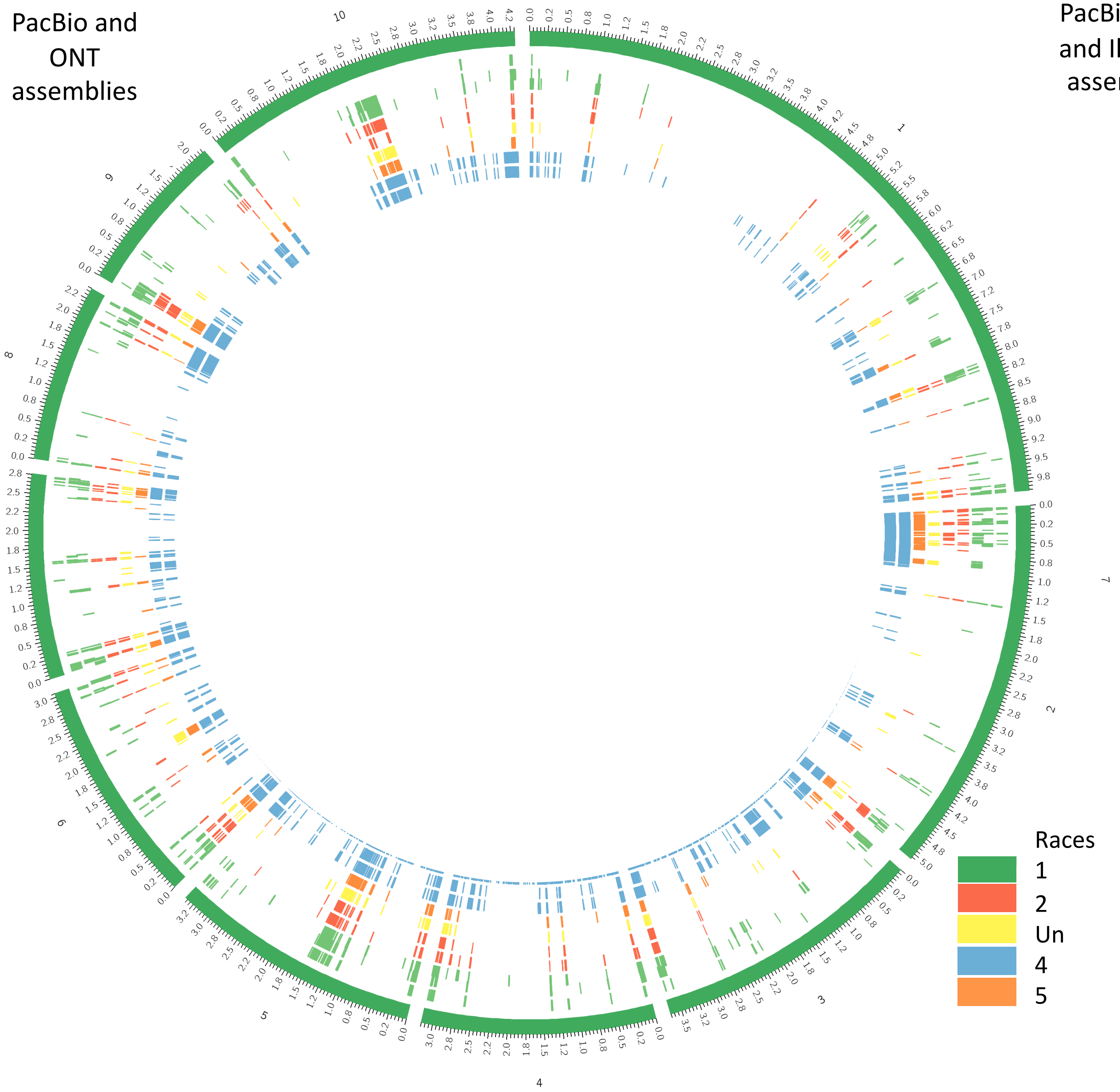

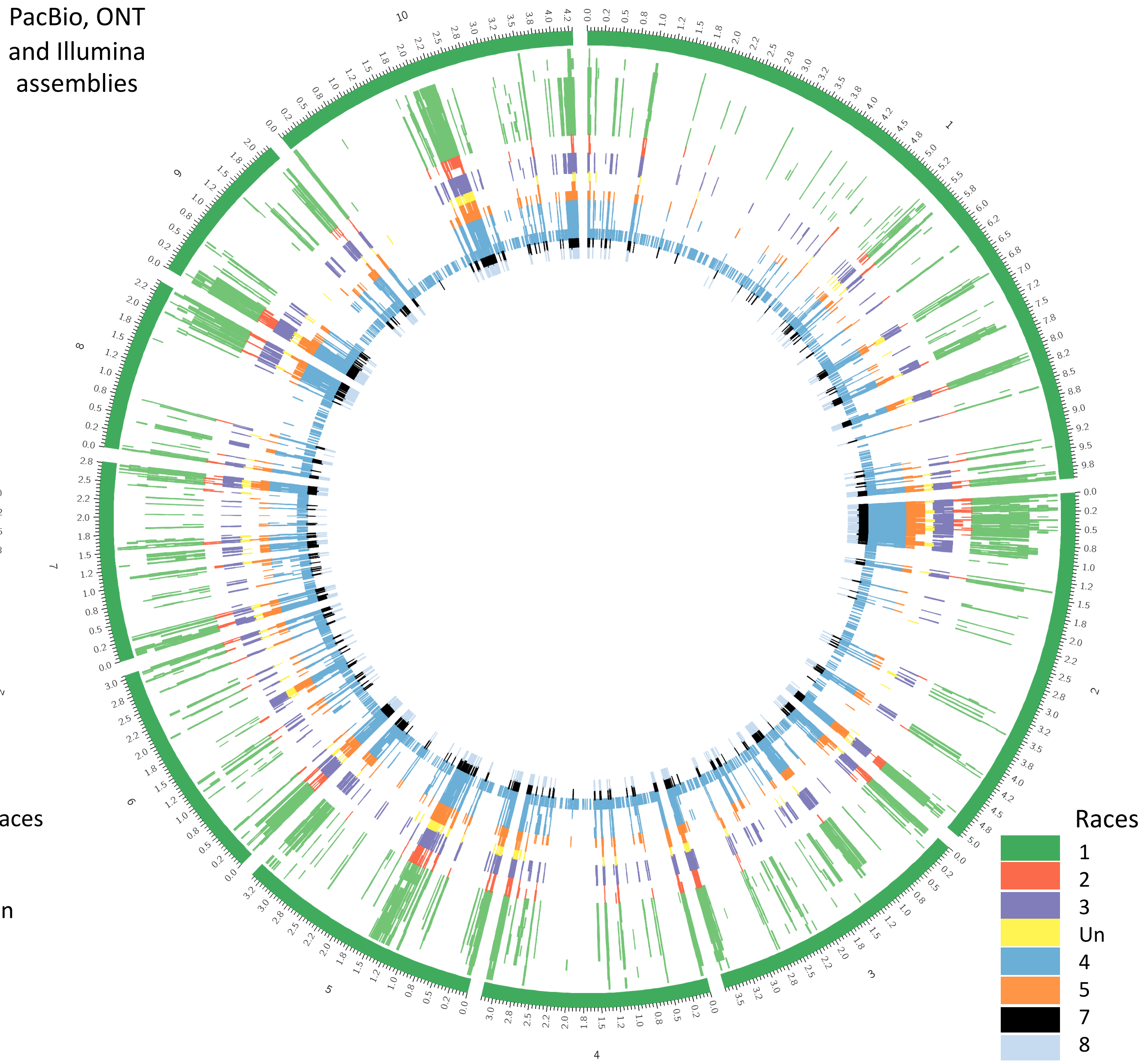

### Supplementary Fig. S11

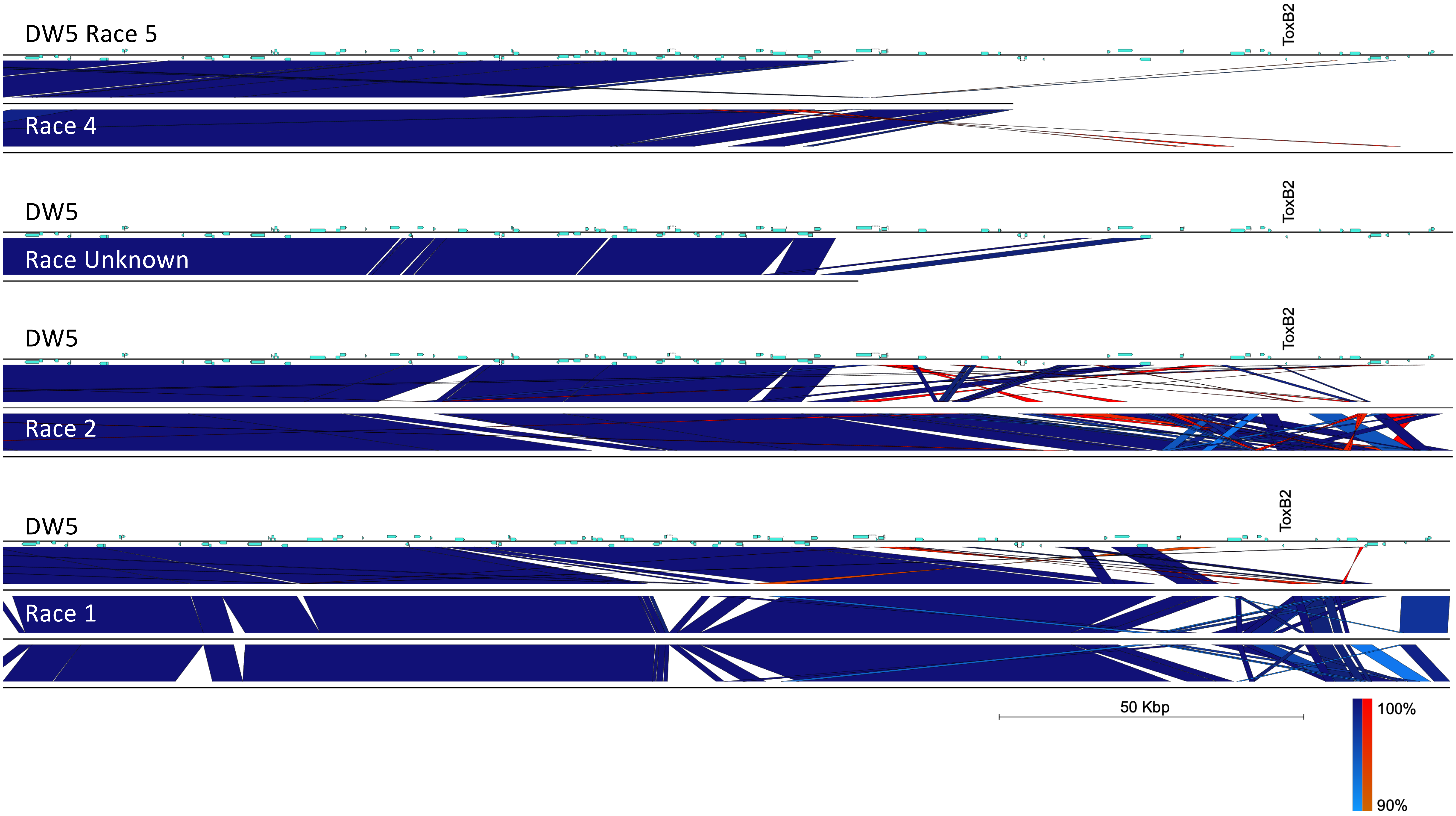

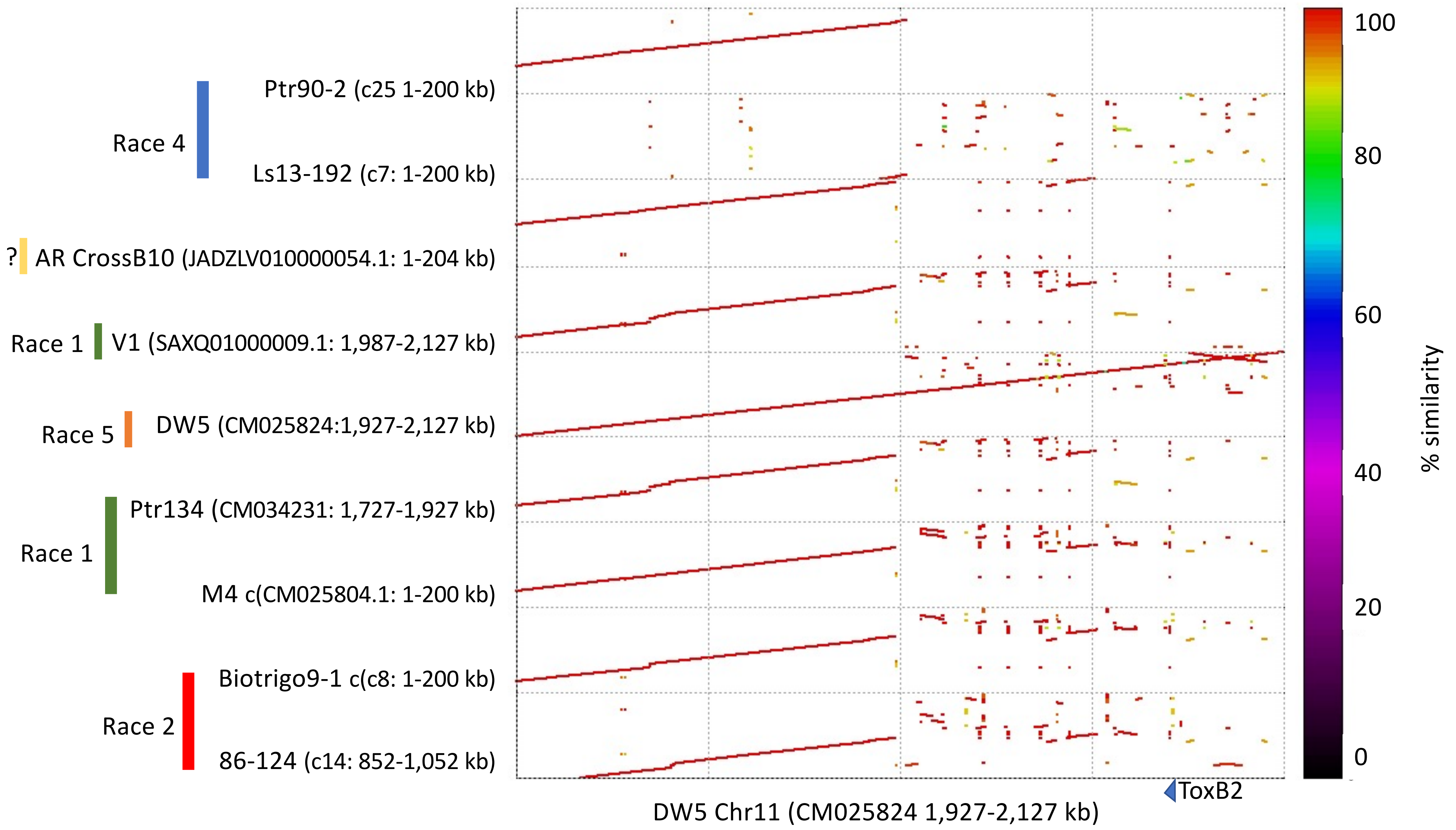

### Supplementary Fig. S12

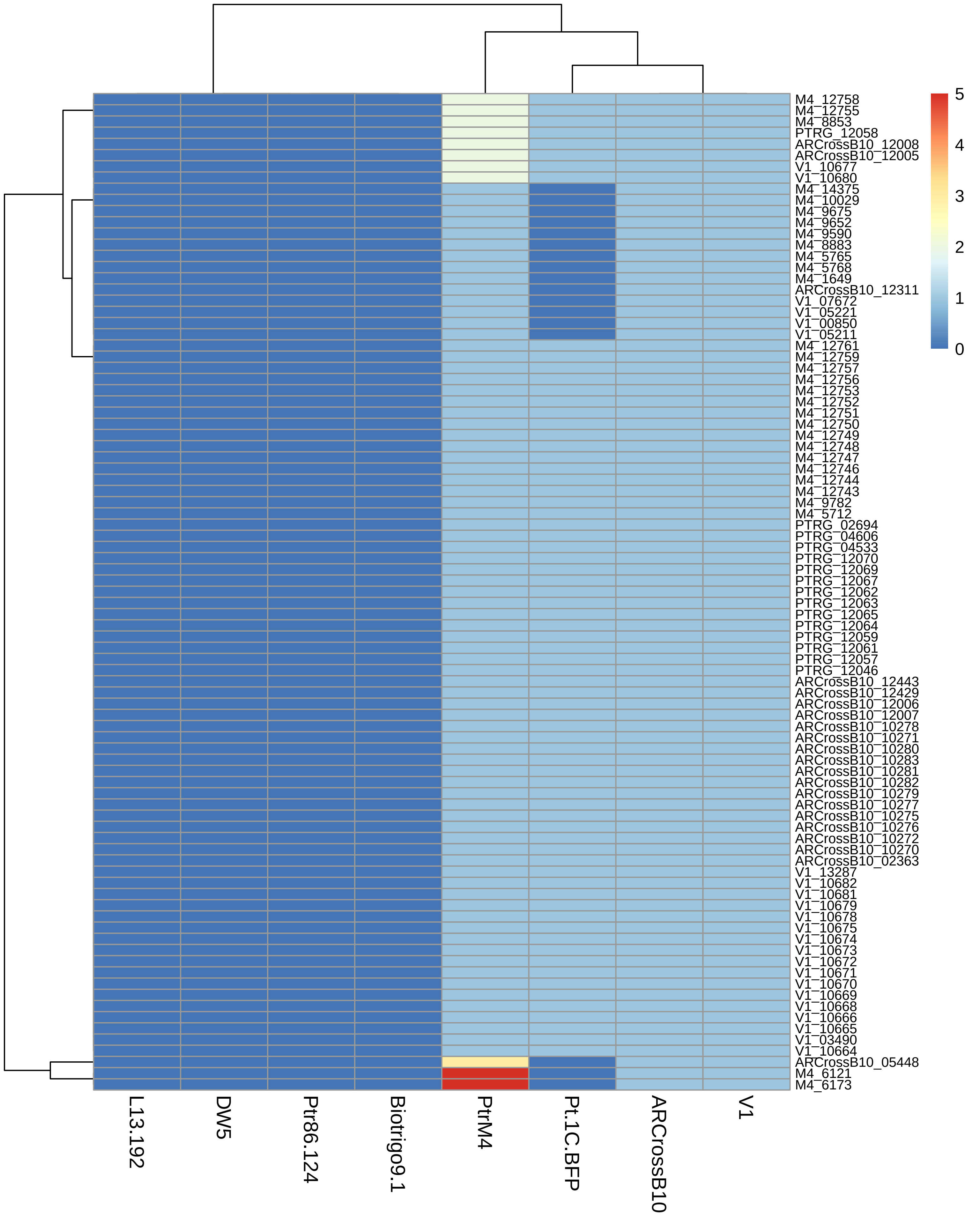

### Supplementary Fig. S13

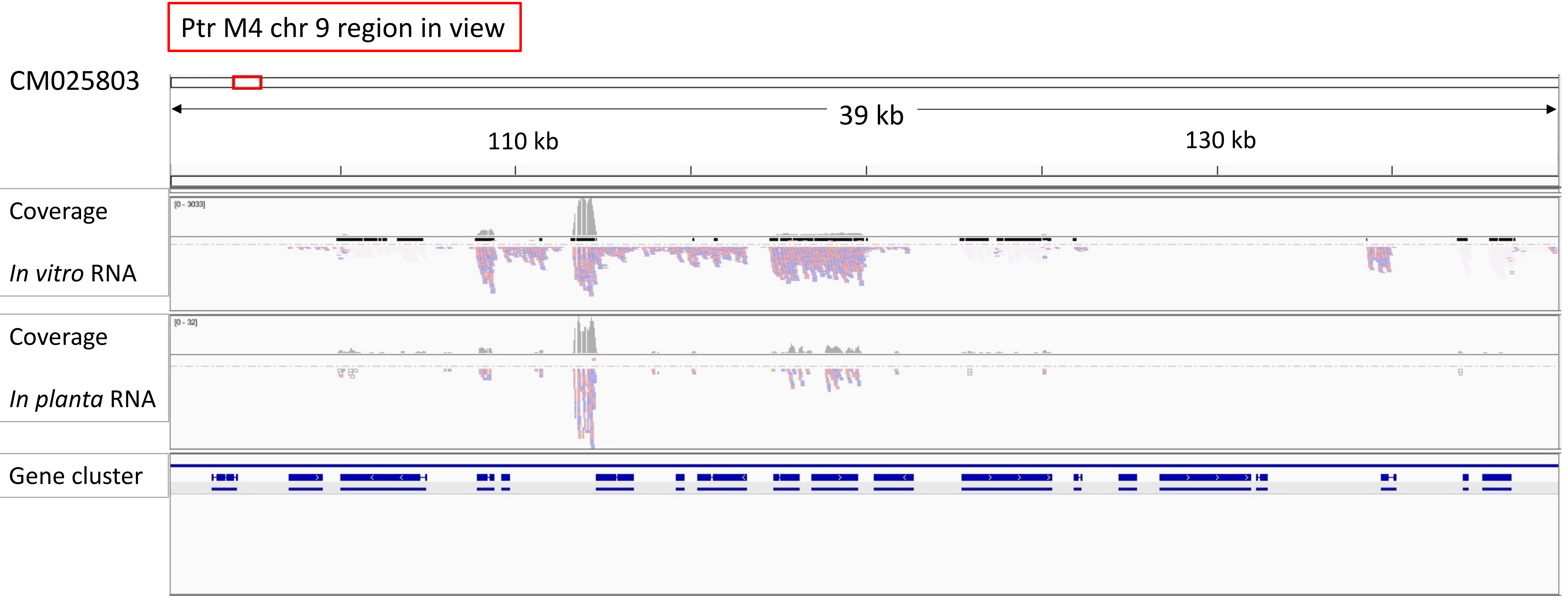

### Supplementary Fig. S14

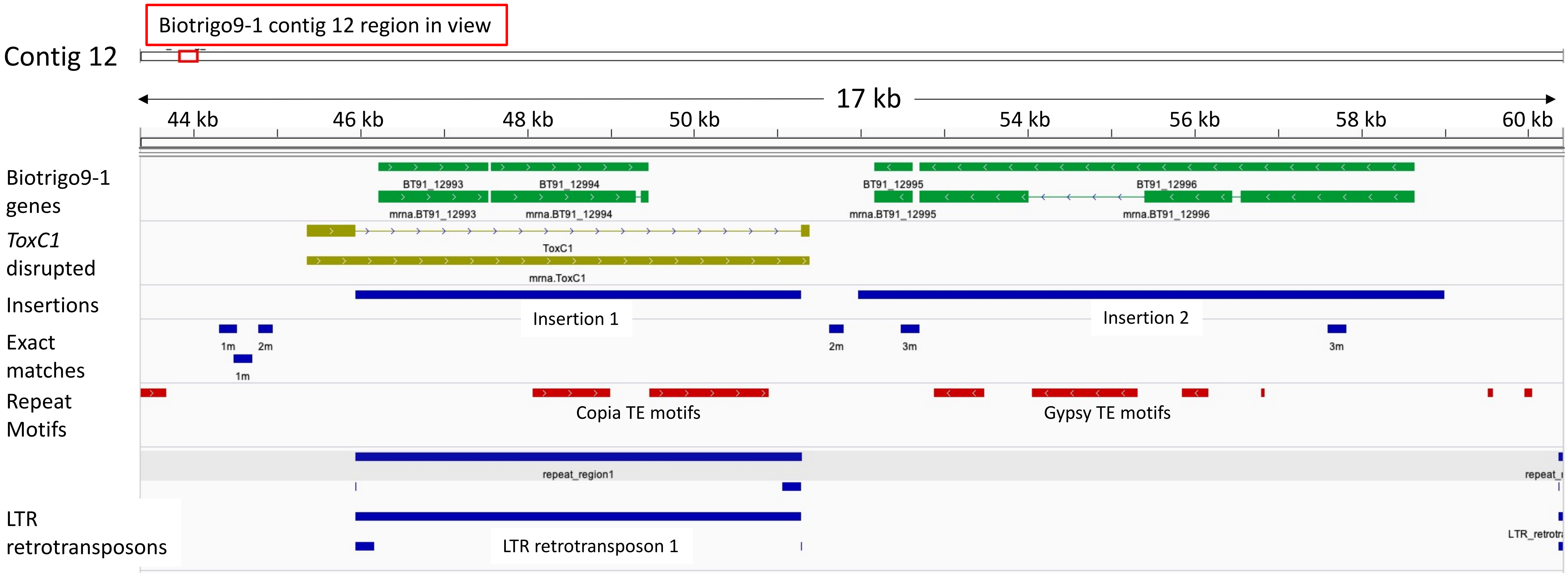

### Supplementary Fig. S15

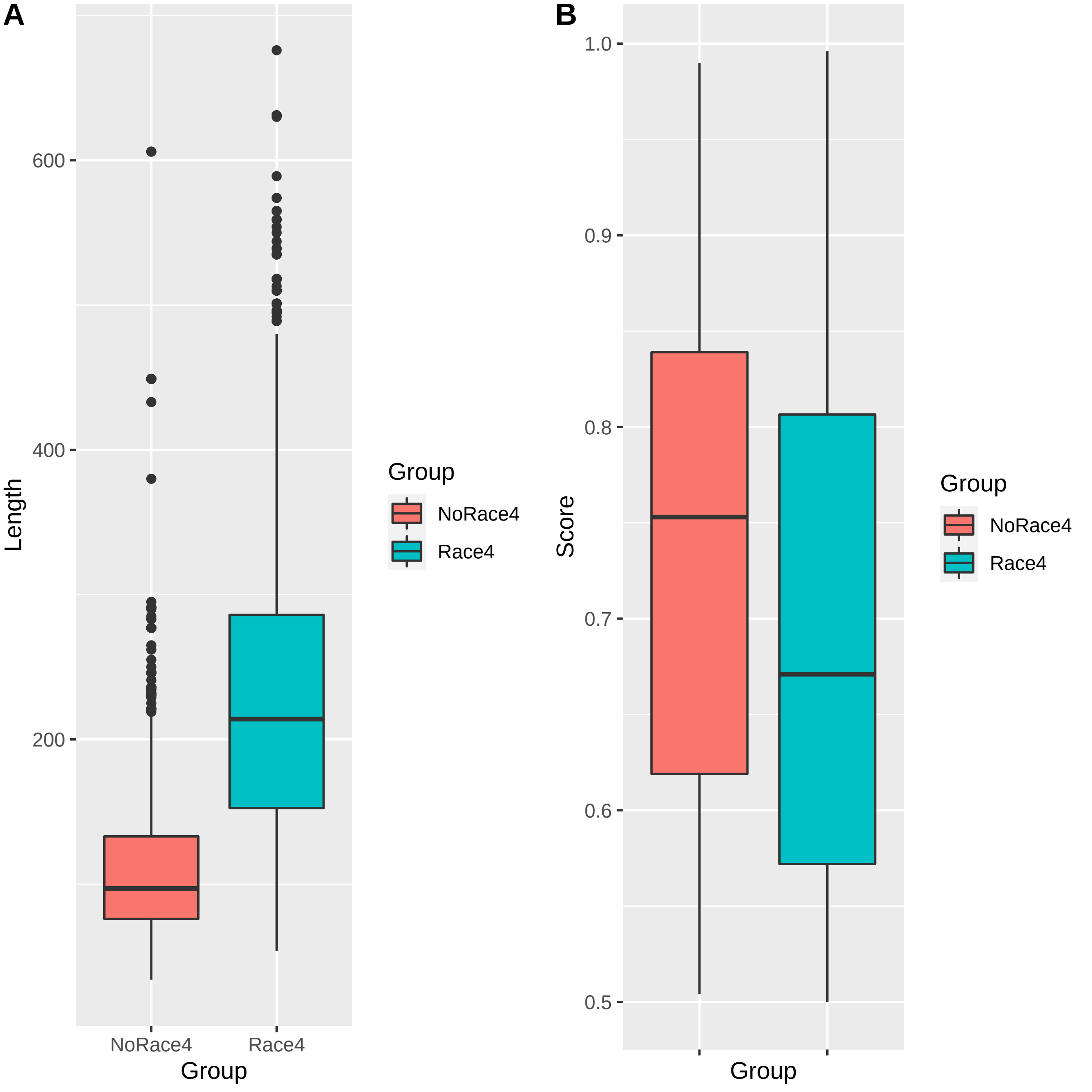
