## Supplementary data 2 for "A global pangenome for the wheat fungal pathogen *Pyrenophora tritici-repentis* and prediction of effector protein structural homology"

```
#Ptr
wget ftp://ftp.ncbi.nlm.nih.gov/genomes/all/GCA/003/231/415/GCA_003231415.2_CUR_PtrDW5_2.1/GCA_003231415.2_CUR_PtrDW5_2.1_genomic.fna.gz
wget ftp://ftp.ncbi.nlm.nih.gov/genomes/all/GCA/003/171/515/GCA_003171515.3_CUR_PTRM4_2.2/GCA_003171515.3_CUR_PTRM4_2.2_genomic.fna.gz
wget ftp://ftp.ncbi.nlm.nih.gov/genomes/all/GCA/003/231/325/GCA_003231325.2_CUR_Ptr134_2.0/GCA_003231325.2_CUR_Ptr134_2.0_genomic.fna.gz
wget ftp://ftp.ncbi.nlm.nih.gov/genomes/all/GCA/008/692/205/GCA_008692205.1_Ptr_V0001/GCA_008692205.1_Ptr_V0001_genomic.fna.gz
wget
ftp://ftp.ncbi.nlm.nih.gov/genomes/all/GCA/018/492/725/GCA_018492725.1_NDSU_Ptr_ARCrossB10_PacBio/GCA_018492725.1_NDSU_Ptr_ARCrossB10_PacBio_gen
omic.fna.gz
wget ftp://ftp.ncbi.nlm.nih.gov/genomes/all/GCA/003/231/425/GCA_003231425.1_Ptr86-124/GCA_003231425.1_Ptr86-124_genomic.fna.gz
wget ftp://ftp.ncbi.nlm.nih.gov/genomes/all/GCA/003/231/345/GCA_003231345.1_Ptr5213/GCA_003231345.1_Ptr5213_genomic.fna.gz
wget ftp://ftp.ncbi.nlm.nih.gov/genomes/all/GCA/003/231/365/GCA_003231365.1_Ptr239/GCA_003231365.1_Ptr239_genomic.fna.gz

#Ptt and Pttm
wget ftp://ftp.ncbi.nlm.nih.gov/genomes/all/GCA/900/232/045/GCA_900232045.3_PTTW11/GCA_900232045.3_PTTW11_protein.faa.gz
wget ftp://ftp.ncbi.nlm.nih.gov/genomes/all/GCA/900/231/935/GCA_900231935.2_ERZ478497/GCA_900231935.2_ERZ478497_protein.faa.gz
wget ftp://ftp.ncbi.nlm.nih.gov/genomes/all/GCA/009/728/675/GCA_009728675.1_ASM972867v1/GCA_009728675.1_ASM972867v1_protein.faa.gz
wget ftp://ftp.ncbi.nlm.nih.gov/genomes/all/GCA/009/728/665/GCA_009728665.1_ASM972866v1/GCA_009728665.1_ASM972866v1_protein.faa.gz
wget ftp://ftp.ncbi.nlm.nih.gov/genomes/all/GCA/009/728/645/GCA_009728645.1_ASM972864v1/GCA_009728645.1_ASM972864v1_protein.faa.gz
wget ftp://ftp.ncbi.nlm.nih.gov/genomes/all/GCA/009/728/635/GCA_009728635.1_ASM972863v1/GCA_009728635.1_ASM972863v1_protein.faa.gz
wget ftp://ftp.ncbi.nlm.nih.gov/genomes/all/GCA/009/728/655/GCA_009728655.1_ASM972865v1/GCA_009728655.1_ASM972865v1_protein.faa.gz
wget ftp://ftp.ncbi.nlm.nih.gov/genomes/all/GCA/000/166/005/GCA_000166005.1_PyrTer_1.0/GCA_000166005.1_PyrTer_1.0_protein.faa.gz


wget ftp://ftp.ncbi.nlm.nih.gov/genomes/all/GCA/020/027/095/GCA_020027095.1_ASM2002709v1/GCA_020027095.1_ASM2002709v1_genomic.fna.gz
wget ftp://ftp.ncbi.nlm.nih.gov/genomes/all/GCA/014/334/815/GCA_014334815.1_ASM1433481v1/GCA_014334815.1_ASM1433481v1_genomic.fna.gz
wget ftp://ftp.ncbi.nlm.nih.gov/genomes/all/GCA/014/334/755/GCA_014334755.1_ASM1433475v1/GCA_014334755.1_ASM1433475v1_genomic.fna.gz
wget ftp://ftp.ncbi.nlm.nih.gov/genomes/all/GCA/014/334/775/GCA_014334775.1_ASM1433477v1/GCA_014334775.1_ASM1433477v1_genomic.fna.gz
wget ftp://ftp.ncbi.nlm.nih.gov/genomes/all/GCA/014/334/795/GCA_014334795.1_ASM1433479v1/GCA_014334795.1_ASM1433479v1_genomic.fna.gz
wget ftp://ftp.ncbi.nlm.nih.gov/genomes/all/GCA/900/231/935/GCA_900231935.2_ERZ478497/GCA_900231935.2_ERZ478497_genomic.fna.gz
wget ftp://ftp.ncbi.nlm.nih.gov/genomes/all/GCA/008/086/845/GCA_008086845.1_ASM808684v1/GCA_008086845.1_ASM808684v1_genomic.fna.gz
wget ftp://ftp.ncbi.nlm.nih.gov/genomes/all/GCA/008/086/725/GCA_008086725.1_ASM808672v1/GCA_008086725.1_ASM808672v1_genomic.fna.gz
wget ftp://ftp.ncbi.nlm.nih.gov/genomes/all/GCA/008/086/755/GCA_008086755.1_ASM808675v1/GCA_008086755.1_ASM808675v1_genomic.fna.gz
wget ftp://ftp.ncbi.nlm.nih.gov/genomes/all/GCA/900/232/045/GCA_900232045.2_ERZ370274/GCA_900232045.2_ERZ370274_genomic.fna.gz
wget ftp://ftp.ncbi.nlm.nih.gov/genomes/all/GCA/008/086/785/GCA_008086785.1_ASM808678v1/GCA_008086785.1_ASM808678v1_genomic.fna.gz
wget ftp://ftp.ncbi.nlm.nih.gov/genomes/all/GCA/009/728/675/GCA_009728675.1_ASM972867v1/GCA_009728675.1_ASM972867v1_genomic.fna.gz
wget ftp://ftp.ncbi.nlm.nih.gov/genomes/all/GCA/009/728/665/GCA_009728665.1_ASM972866v1/GCA_009728665.1_ASM972866v1_genomic.fna.gz
wget ftp://ftp.ncbi.nlm.nih.gov/genomes/all/GCA/009/728/645/GCA_009728645.1_ASM972864v1/GCA_009728645.1_ASM972864v1_genomic.fna.gz
wget ftp://ftp.ncbi.nlm.nih.gov/genomes/all/GCA/009/728/635/GCA_009728635.1_ASM972863v1/GCA_009728635.1_ASM972863v1_genomic.fna.gz
wget ftp://ftp.ncbi.nlm.nih.gov/genomes/all/GCA/009/728/655/GCA_009728655.1_ASM972865v1/GCA_009728655.1_ASM972865v1_genomic.fna.gz
wget ftp://ftp.ncbi.nlm.nih.gov/genomes/all/GCA/006/112/615/GCA_006112615.1_ASM611261v1/GCA_006112615.1_ASM611261v1_genomic.fna.gz

#Ps
wget https://ftp.ncbi.nlm.nih.gov/genomes/all/GCA/000/465/215/GCA_000465215.2_Pysem1.0/GCA_000465215.2_Pysem1.0_genomic.fna.gz

#Pg
wget https://ftp.ncbi.nlm.nih.gov/genomes/all/GCA/012/365/135/GCA_012365135.1_ASM1236513v1/GCA_012365135.1_ASM1236513v1_genomic.fna.gz


#Bm
wget https://ftp.ncbi.nlm.nih.gov/genomes/all/GCF/000/354/255/GCF_000354255.1_CocheC4_1/GCF_000354255.1_CocheC4_1_genomic.fna.gz
```

```
#Bv
wget
https://ftp.ncbi.nlm.nih.gov/genomes/all/GCF/000/527/765/GCF_000527765.1_Cochliobolus_victoriae_v1.0/GCF_000527765.1_Cochliobolus_victoriae_v1.0
_genomic.fna.gz

#Bz
wget ftp://ftp.ncbi.nlm.nih.gov/genomes/all/GCA/016/906/865/GCA_016906865.1_ASM1690686v1

#Bo
wget
https://ftp.ncbi.nlm.nih.gov/genomes/all/GCF/000/523/455/GCF_000523455.1_Cochliobolus_miyabeanus_v1.0/GCF_000523455.1_Cochliobolus_miyabeanus_v1
.0_genomic.fna.gz

#Bs
wget ftp://ftp.ncbi.nlm.nih.gov/genomes/all/GCA/008/452/705/GCA_008452705.1_ASM845270v1/GCA_008452705.1_ASM845270v1_genomic.fna.gz
wget ftp://ftp.ncbi.nlm.nih.gov/genomes/all/GCA/008/452/735/GCA_008452735.1_ASM845273v1/GCA_008452735.1_ASM845273v1_genomic.fna.gz
wget ftp://ftp.ncbi.nlm.nih.gov/genomes/all/GCA/008/452/725/GCA_008452725.1_ASM845272v1/GCA_008452725.1_ASM845272v1_genomic.fna.gz
wget ftp://ftp.ncbi.nlm.nih.gov/genomes/all/GCA/008/452/715/GCA_008452715.1_ASM845271v1/GCA_008452715.1_ASM845271v1_genomic.fna.gz
wget ftp://ftp.ncbi.nlm.nih.gov/genomes/all/GCA/004/329/375/GCA_004329375.1_ASM432937v1/GCA_004329375.1_ASM432937v1_genomic.fna.gz
wget ftp://ftp.ncbi.nlm.nih.gov/genomes/all/GCA/013/416/765/GCA_013416765.1_ASM1341676v1/GCA_013416765.1_ASM1341676v1_genomic.fna.gz
wget ftp://ftp.ncbi.nlm.nih.gov/genomes/all/GCA/000/338/995/GCA_000338995.1_Cocsa1/GCA_000338995.1_Cocsa1_genomic.fna.gz

#Bc
wget https://ftp.ncbi.nlm.nih.gov/genomes/all/GCA/002/286/855/GCA_002286855.1_ASM228685v1/GCA_002286855.1_ASM228685v1_genomic.fna.gz


#Ssp.
wget https://ftp.ncbi.nlm.nih.gov/genomes/all/GCA/001/644/525/GCA_001644525.1_Stasp1/GCA_001644525.1_Stasp1_genomic.fna.gz

#Pa
wget ftp://ftp.ncbi.nlm.nih.gov/genomes/all/GCA/003/501/935/GCA_003501935.1_ASM350193v1/GCA_003501935.1_ASM350193v1_genomic.fna.gz
wget ftp://ftp.ncbi.nlm.nih.gov/genomes/all/GCA/003/501/955/GCA_003501955.1_ASM350195v1/GCA_003501955.1_ASM350195v1_genomic.fna.gz
wget ftp://ftp.ncbi.nlm.nih.gov/genomes/all/GCA/003/503/115/GCA_003503115.1_ASM350311v1/GCA_003503115.1_ASM350311v1_genomic.fna.gz
wget ftp://ftp.ncbi.nlm.nih.gov/genomes/all/GCA/003/503/205/GCA_003503205.1_ASM350320v1/GCA_003503205.1_ASM350320v1_genomic.fna.gz
wget ftp://ftp.ncbi.nlm.nih.gov/genomes/all/GCA/003/503/125/GCA_003503125.1_ASM350312v1/GCA_003503125.1_ASM350312v1_genomic.fna.gz
wget ftp://ftp.ncbi.nlm.nih.gov/genomes/all/GCA/003/501/975/GCA_003501975.1_ASM350197v1/GCA_003501975.1_ASM350197v1_genomic.fna.gz
wget ftp://ftp.ncbi.nlm.nih.gov/genomes/all/GCA/003/502/015/GCA_003502015.1_ASM350201v1/GCA_003502015.1_ASM350201v1_genomic.fna.gz
wget ftp://ftp.ncbi.nlm.nih.gov/genomes/all/GCA/003/502/495/GCA_003502495.1_ASM350249v1/GCA_003502495.1_ASM350249v1_genomic.fna.gz
wget ftp://ftp.ncbi.nlm.nih.gov/genomes/all/GCA/003/503/195/GCA_003503195.1_ASM350319v1/GCA_003503195.1_ASM350319v1_genomic.fna.gz
wget ftp://ftp.ncbi.nlm.nih.gov/genomes/all/GCA/003/503/165/GCA_003503165.1_ASM350316v1/GCA_003503165.1_ASM350316v1_genomic.fna.gz

#Pn
wget ftp://ftp.ncbi.nlm.nih.gov/genomes/all/GCA/002/267/025/GCA_002267025.1_ASM226702v1/GCA_002267025.1_ASM226702v1_genomic.fna.gz
wget ftp://ftp.ncbi.nlm.nih.gov/genomes/all/GCA/002/267/045/GCA_002267045.1_ASM226704v1/GCA_002267045.1_ASM226704v1_genomic.fna.gz
wget ftp://ftp.ncbi.nlm.nih.gov/genomes/all/GCA/002/267/005/GCA_002267005.1_ASM226700v1/GCA_002267005.1_ASM226700v1_genomic.fna.gz
wget ftp://ftp.ncbi.nlm.nih.gov/genomes/all/GCA/008/452/785/GCA_008452785.1_ASM845278v1/GCA_008452785.1_ASM845278v1_genomic.fna.gz
wget ftp://ftp.ncbi.nlm.nih.gov/genomes/all/GCA/016/801/405/GCA_016801405.1_ASM1680140v1/GCA_016801405.1_ASM1680140v1_genomic.fna.gz


#Zt
wget ftp://ftp.ncbi.nlm.nih.gov/genomes/all/GCA/002/937/425/GCA_002937425.1_ASM293742v1/GCA_002937425.1_ASM293742v1_genomic.fna.gz
```

```
wget ftp://ftp.ncbi.nlm.nih.gov/genomes/all/GCA/017/562/065/GCA_017562065.1_ASM1756206v1/GCA_017562065.1_ASM1756206v1_genomic.fna.gz
wget ftp://ftp.ncbi.nlm.nih.gov/genomes/all/GCA/902/712/725/GCA_902712725.1_ERS3651594/GCA_902712725.1_ERS3651594_genomic.fna.gz
wget ftp://ftp.ncbi.nlm.nih.gov/genomes/all/GCA/017/562/105/GCA_017562105.1_ASM1756210v1/GCA_017562105.1_ASM1756210v1_genomic.fna.gz
wget ftp://ftp.ncbi.nlm.nih.gov/genomes/all/GCA/017/766/645/GCA_017766645.1_ASM1776664v1/GCA_017766645.1_ASM1776664v1_genomic.fna.gz
wget ftp://ftp.ncbi.nlm.nih.gov/genomes/all/GCA/017/766/825/GCA_017766825.1_ASM1776682v1/GCA_017766825.1_ASM1776682v1_genomic.fna.gz
wget ftp://ftp.ncbi.nlm.nih.gov/genomes/all/GCA/019/443/865/GCA_019443865.1_ASM1944386v1/GCA_019443865.1_ASM1944386v1_genomic.fna.gz
wget ftp://ftp.ncbi.nlm.nih.gov/genomes/all/GCA/017/766/805/GCA_017766805.1_ASM1776680v1/GCA_017766805.1_ASM1776680v1_genomic.fna.gz
wget ftp://ftp.ncbi.nlm.nih.gov/genomes/all/GCA/019/443/905/GCA_019443905.1_ASM1944390v1/GCA_019443905.1_ASM1944390v1_genomic.fna.gz
wget ftp://ftp.ncbi.nlm.nih.gov/genomes/all/GCA/017/766/425/GCA_017766425.1_ASM1776642v1/GCA_017766425.1_ASM1776642v1_genomic.fna.gz
wget ftp://ftp.ncbi.nlm.nih.gov/genomes/all/GCA/017/766/785/GCA_017766785.1_ASM1776678v1/GCA_017766785.1_ASM1776678v1_genomic.fna.gz
wget ftp://ftp.ncbi.nlm.nih.gov/genomes/all/GCA/017/766/145/GCA_017766145.1_ASM1776614v1/GCA_017766145.1_ASM1776614v1_genomic.fna.gz
wget ftp://ftp.ncbi.nlm.nih.gov/genomes/all/GCA/017/766/525/GCA_017766525.1_ASM1776652v1/GCA_017766525.1_ASM1776652v1_genomic.fna.gz
wget ftp://ftp.ncbi.nlm.nih.gov/genomes/all/GCA/019/443/885/GCA_019443885.1_ASM1944388v1/GCA_019443885.1_ASM1944388v1_genomic.fna.gz
wget ftp://ftp.ncbi.nlm.nih.gov/genomes/all/GCA/017/766/145/GCA_017766145.1_ASM1776614v1/GCA_017766145.1_ASM1776614v1_genomic.fna.gz
wget ftp://ftp.ncbi.nlm.nih.gov/genomes/all/GCA/017/766/525/GCA_017766525.1_ASM1776652v1/GCA_017766525.1_ASM1776652v1_genomic.fna.gz
wget ftp://ftp.ncbi.nlm.nih.gov/genomes/all/GCA/017/766/725/GCA_017766725.1_ASM1776672v1/GCA_017766725.1_ASM1776672v1_genomic.fna.gz
wget ftp://ftp.ncbi.nlm.nih.gov/genomes/all/GCA/000/219/625/GCA_000219625.1_MYCGR_v2.0/GCA_000219625.1_MYCGR_v2.0_genomic.fna.gz
wget
ftp://ftp.ncbi.nlm.nih.gov/genomes/all/GCA/900/099/495/GCA_900099495.1_ZT1A5_complete_assembly/GCA_900099495.1_ZT1A5_complete_assembly_genomic.f
na.gz
wget ftp://ftp.ncbi.nlm.nih.gov/genomes/all/GCA/900/184/115/GCA_900184115.1_ST99CH_1E4/GCA_900184115.1_ST99CH_1E4_genomic.fna.gz
wget ftp://ftp.ncbi.nlm.nih.gov/genomes/all/GCA/900/184/105/GCA_900184105.1_ST99CH_3D1/GCA_900184105.1_ST99CH_3D1_genomic.fna.gz
wget ftp://ftp.ncbi.nlm.nih.gov/genomes/all/GCA/900/091/695/GCA_900091695.1_Zt_ST99CH_3D7/GCA_900091695.1_Zt_ST99CH_3D7_genomic.fna.gz
wget ftp://ftp.ncbi.nlm.nih.gov/genomes/all/GCA/002/937/415/GCA_002937415.1_ASM293741v1/GCA_002937415.1_ASM293741v1_genomic.fna.gz
wget ftp://ftp.ncbi.nlm.nih.gov/genomes/all/GCA/000/223/645/GCA_000223645.2_ASM22364v2/GCA_000223645.2_ASM22364v2_genomic.fna.gz
```
